# Chemotherapy-Induced Oral Mucosal Injury Is Defined by p53 Activation, Cell Cycle Arrest and Diverse Epithelial Progenitor Dynamics

**DOI:** 10.64898/2026.04.06.716752

**Authors:** Pedro H. F. Silva, Lu Li, Madison Muriel, Tara Brennan, Md Hasanuzzaman, Mariadela Demoura, Joleene Stampfer, Michele Sveinsson, Christos Fountzilas, Jill E. Schabert, Anurag K. Singh, Jonathan E. Bard, Nicolas F. Schlecht, Laertis Ikonomou, Patricia I. Diaz

**Affiliations:** Department of Oral Biology, University at Buffalo, The State University of New York, Buffalo, New York, USA; Department of Cancer Prevention & Control, Roswell Park Comprehensive Cancer Center, Buffalo, New York, USA; Department of Medicine, Roswell Park Comprehensive Cancer Center, Buffalo, New York, USA; Department of Dentistry and Maxillofacial Prosthetics, Roswell Park Comprehensive Cancer Center, Buffalo, New York, USA; Department of Radiation Medicine, Roswell Park Comprehensive Cancer Center, Buffalo, New York, USA; Department of Biochemistry, University at Buffalo, The State University of New York, Buffalo New York, USA; Division of Pulmonary, Critical Care and Sleep Medicine, Department of Medicine, University at Buffalo, The State University of New York, Buffalo, New ork, USA

## Abstract

Chemotherapy-induced oral mucositis is a common and debilitating complication, yet the mechanisms underlying oral mucosa injury and repair remain poorly defined. Using a mouse model of 5-fluorouracil (5-FU)–induced mucositis, we define gene networks and oral mucosal cellular landscape dynamics in response to chemotoxic stress. We show that 5-FU–induced epithelial atrophy is driven primarily by cell cycle arrest rather than apoptosis, despite concurrent activation of p53-dependent transcriptional programs linked to both cell fates within individual cells. Relative to intestine, the recovering oral mucosa exhibits a more effective cell cycle checkpoint response and uniquely undergoes metabolic reprogramming toward lipid oxidation. Single-cell RNA sequencing revealed putative epithelial progenitor populations with distinct responses to chemotherapy, including chemoresistant cells with a p53- and AP-1 complex gene signature, reminiscent of lung transitional cell states. These findings define diverse progenitor dynamics and p53-driven responses as key determinants of oral mucosal injury and repair following chemotherapy.

## Introduction

The integrity of the oral and gastrointestinal mucosae is essential for preserving the various functions of these barrier tissues, which include nutrient acquisition, prevention of microbial invasion, and regulation of local and systemic immune homeostasis. However, the way these mucosae maintain health and recover from different injuries is still incompletely understood. This is especially true for the oral mucosa, which is considered a paradigm of rapid wound healing. The fast recovery from injury of the oral barrier is attributed to having epithelial progenitor populations with higher regenerative capacity compared to other tissues with slower healing potential, such as the skin^1^. Nevertheless, the cellular populations and gene regulatory networks in oral tissues mediating regeneration after injury are undefined. Understanding responses of upper (oral) and lower gastrointestinal mucosae to various insults can inform the development of therapies against diseases affecting these barriers.

One type of insult that affects gastrointestinal mucosal compartments is chemotoxic stress secondary to radiation and chemotherapy for cancer. The main clinical manifestation of chemotoxic injury is gastrointestinal mucositis, a condition characterized by erythema and ulcer-like lesions in the oral cavity and intestine, sometimes accompanied by diarrhea^2–4^. Although mucositis self-resolves within days or weeks of presentation, it can cause significant pain, treatment interruptions and increase the risk of bloodstream infections^5–7^. Human histological evidence and animal models show mucositis is characterized by epithelial atrophy due to impairment of tissue self-renewal^8–10^. In the intestinal mucosa, chemotherapy triggers a modest and temporary increase in apoptosis of crypt epithelial cells, accompanied by a long-lasting decrease in their proliferation^10,11^. Conceptual models of oral mucositis pathophysiology also point to apoptosis of basal epithelial cells, in addition to activation of pro-inflammatory pathways, such as Nuclear factor-kappa B signaling, as mediators of oral mucosal injury^12,13^. There is, however, scant evidence from humans or animal models outlining the processes involved in chemotoxic injury of oral mucosal tissues, and it is unknown whether oral and intestinal tissues respond to chemotherapy in the same manner.

The sensitivity of cancer and non-cancer cells to chemotherapeutic agents is highly dependent on the activity of the transcription factor p53 which is involved in regulation of cell responses to DNA damage by inducing cell death, arresting replication, and facilitating survival^10,14,15^. In vivo experiments have shown that the response in intestinal tissues to the commonly used chemotherapeutic 5-fluorouracil (5-FU) is dependent on p53^10^. However, the effects of p53 activation in response to DNA-damaging agents appear to be distinct across tissues. For instance, while in the hematopoietic compartment p53 is responsible for cell apoptosis and ablation in response to radiation, causing lethality^16^, p53 has a protective effect in the intestinal mucosa decreasing the severity of radiation-induced mucositis and survival by activating cell cycle checkpoints^17–19^. While it is expected that the response to chemotherapeutic drugs in the oral mucosa will also be mediated by p53, the nature of the transcriptional program activated in oral tissues by chemotherapeutic drugs is unknown and it could differ from that in the intestine as p53 activates distinct transcriptional targets in a cell- and tissue-specific manner^18–20^. Systematic in vivo studies are needed to help understand the kinetics of the transcriptional responses to chemotherapeutic agents in different gastrointestinal mucosal tissues that can help identify actionable therapeutic targets for mucositis.

Here we use a mouse model to examine the responses of the oral and small intestinal mucosae during mucositis development after 5-FU administration. Our work shows that although a p53-driven transcriptional response dominates in oral and intestinal mucosa, regulation of p53 targets is tissue-specific. We also show that although p53 target genes involved in controlling divergent cell fates were co-expressed within tissues and within single cells in response to 5-FU, epithelial cell cycle arrest, and not apoptosis, was the hallmark event associated with mucositis development in both oral and intestinal mucosae. In the oral tissues, however, we identified metabolic reprogramming towards fatty acid oxidation as a site-specific response potentially contributing to tissue recovery. Through single cell RNA Sequencing (scRNASeq) of oral mucosa we confirmed that epithelial basal proliferative cells are highly susceptible to the effects of 5-FU, together with a population of Wnt signaling-responsive putative oral epithelial cell progenitors. We also uncovered an epithelial progenitor-like population with a p53 and Adaptor Protein-1 (AP-1) complex gene signature that appeared resistant to the effects of 5-FU. This population has transcriptional similarities to transitional cells involved in regeneration in the respiratory tract^21–23^ and emerges as a candidate population that mediates recovery from chemotoxic injury in the oral mucosa.

## Methods

### Mucositis animal model

Eight-week-old C57BL/6 mice (The Jackson Laboratory, Bar Harbor, ME, USA) were used for all experiments. All animal studies were conducted in accordance with the Animal Welfare Act and the recommendations outlined in the Guide for the Care and Use of Laboratory Animals (National Institutes of Health). All experimental protocols were reviewed and approved by the Institutional Animal Care and Use Committee (IACUC) at the University at Buffalo (Protocol 202000086).

5-Fluorouracil (5-FU) powder (Sigma-Aldrich, St. Louis, MO, USA; Cat# F6627) was prepared by dissolving it in sterile 0.85% saline to a final concentration of 6.25 mg ml^-1^. The solution was incubated at 60°C for 45 minutes with periodic vortexing to achieve complete dissolution. The pH was then adjusted to 9.2. Prepared samples were aliquoted, stored at −20°C, and each aliquot was thawed only once before being administered to mice.

Mice received 5-FU via intraperitoneal injection at 100 mg/kg at days 0 and 1, followed by 50 mg/kg daily for 5 more days. Animals were monitored until day 10. Body weight was recorded daily. Mice were sacrificed at days 1, 2, 4, 6, 8, and 10 and tongue and small intestine tissue samples were collected. Control mice received PBS injections according to the same schedule. For single cell RNA sequencing (scRNASeq) experiments, the 5-FU administration scheme was shortened following the same procedures but stopping 5-FU administration after day 4 and mice were sacrificed at days 2, 4 and 8.

### Toluidine blue staining and quantification of tongue lesions

Macroscopic lesions in tongues were assessed using toluidine blue staining. Freshly excised tongues were gently rinsed in phosphate-buffered saline (PBS; Corning, Corning, NY, USA; Cat# 46-013-CM) and then immersed in 1% (w/v) toluidine blue (Sigma-Aldrich, St. Louis, MO, USA; Cat# T3260) prepared in 1% (v/v) acetic acid (Sigma-Aldrich, St. Louis, MO, USA; Cat# 695092) for 2–3 min at room temperature. Excess dye was removed by brief rinsing in 1% acetic acid followed by rinsing in PBS. Stained tissues were photographed using a Canon DSLR camera (Canon Inc., Tokyo, Japan) under standardized lighting conditions. Regions retaining toluidine blue staining were considered indicative of epithelial breaches. Lesion burden was quantified using ImageJ software (NIH, Bethesda, MD, USA) by measuring the total toluidine blue–positive area and normalizing it to the total tongue area for each sample. Data are presented as the percentage of tongue surface area exhibiting positive toluidine blue staining.

### Histological analyses

Tongue and intestinal tissues were collected at designated experimental time points following treatment. Intestinal tissues were flushed with PBS. Tissues were then immediately fixed in 4% paraformaldehyde (Electron Microscopy Sciences, Hatfield, PA, USA; Cat# 15710), dehydrated with a series of ethanol washes, embedded in paraffin (Fisher Scientific, Waltham, MA, USA; Cat# P31-500), and sectioned. Five μm sections were stained with hematoxylin and eosin (H&E) using standard protocols. Micrographs were acquired using a Nikon light microscope (Model Eclipse Ni-U, 944389, Nikon Instruments Inc., Melville, NY, USA) equipped with a PCO.edge camera (Excelitas PCO GmbH, Kelheim, Germany). Histomorphometric analysis was performed using NIS-Elements AR software (version 6.10.02; Nikon Instruments Inc., Melville, NY, USA). For tongue, epithelial injury was quantified by measuring epithelial area (μm²) normalized to a defined linear length of the dorsal epithelium. In addition, basal cell density was assessed by counting the number of basal epithelial cells within a standardized 500 μm length of the epithelial layer. In some tongues, macroscopic lesions visualized with toluidine blue were tracked in tissue sections by placing a 6.0 silk suture next to the lesion. For intestine, mucosal injury was evaluated by measuring the length of villi, defined as the distance from the villus tip to the villus–crypt junction, and calculated as the average length of 10 individual villi per section. Multiple non-overlapping fields of view were analyzed per tissue section, and values were averaged to obtain a representative measurement for each sample.

### Immunohistochemistry

Tongue and small intestine paraffin-embedded tissue sections were cooled to room temperature and added to a Leica Biosystems Bond Rx automated stainer, where they were deparaffinized with Bond Dewax Solution (Leica Biosystems, Buffalo Grove, IL, USA; Cat# AR9222) and rinsed in water. Bond Epitope Retrieval 1 (Leica Biosystems, Buffalo Grove, IL, USA; Cat# AR9961) was used for target retrieval for 20 minutes for CD45 staining. Bond Epitope Retrieval 2 (Leica Biosystems, Buffalo Grove, IL, USA; Cat# AR9640) was used for 30 min for KI67 and cleaved Caspase 3 staining. Slides were blocked using peroxide block from Bond Polymer Refine Detection kit (Leica Biosystems, Buffalo Grove, IL, USA; Cat# DS9800) for 5 minutes. Slides were incubated with an anti-CD45 antibody (rat anti-mouse CD45, BioLegend, San Diego, CA, USA; Cat# 103102, RRID: AB_312967) at 1:30 for 20 min, followed by goat anti-rat (BD Biosciences, San Jose, CA, USA; Cat# 554014, RRID: AB_395289) at 1:100 for 30 min, followed by VECTASTAIN Elite ABC-HRP (Vector Laboratories, Burlingame, CA, USA; Cat# PK6100) for 30 min. Slides were incubated with an anti-cleaved Caspase-3 (Asp175) antibody (Cell Signaling Technology, Danvers, MA, USA; Cat# 9661, RRID:AB_2341188) at 1:150 or an anti-KI67 antibody (R&D Systems, Minneapolis, MN, USA; Cat# MAB7617) at 1:125 for 20 min followed by Rabbit Envision (Dako, Carpinteria, CA, USA; Cat# K4003) for 30 min. Diaminobenzidine (DAB) was applied for 10 minutes for visualization. Slides were counterstained with hematoxylin for 8 minutes and then placed into water. Both DAB and hematoxylin were from the Bond Polymer Refine Detection kit (Leica Biosystems, Buffalo Grove, IL, USA; Cat# DS9800). After removing slides from the Leica Bond Rx they were dehydrated, cleared, and mounted.

Brightfield micrographs were captured using a Nikon microscope (Model Eclipse Ni-U, 944389, Nikon Instruments Inc., Melville, NY, USA) and analyzed using NIS-Elements AR software (version 6.10.02; Nikon Instruments Inc., Melville, NY, USA). CD45-, KI67- and cleaved Caspase-3-positive cells were quantified in defined regions of interest. In tongue sections, positive cells were counted within an epithelial region spanning 500 μm in length. In small intestine sections, positive cells were quantified within a 5000 μm² area in well-oriented crypt–villus units. Multiple fields per section were analyzed, and quantification was performed blinded to treatment group.

### Terminal deoxynucleotidyl transferase dUTP nick end labeling (TUNEL) assays

Cell death associated with DNA strand breaks was assessed using the Click-iT™ Plus TUNEL Assay (Thermo Fisher Scientific, Waltham, MA, USA; Cat# C10617) according to the manufacturer’s instructions with minor modifications. Samples were washed and permeabilized using Proteinase K (Thermo Fisher Scientific, Waltham, MA, USA; Cat# 25530049) diluted 1:25 in PBS for 15 min, followed by refixation. DNA strand breaks were labeled by incubation with terminal deoxynucleotidyl transferase (TdT) reaction buffer and TdT enzyme mixture at 37°C. Fluorescent labeling was achieved using Click-iT™ chemistry, followed by washes in PBS containing 3% bovine serum albumin (BSA; Sigma-Aldrich, St. Louis, MO, USA; Cat# A7906) and 0.1% Triton X-100 (Sigma-Aldrich, St. Louis, MO, USA; Cat# T8787). Nuclei were counterstained with 4′,6-diamidino-2-phenylindole (DAPI; Sigma-Aldrich, St. Louis, MO, USA; Cat# D9542) for 15 min at room temperature. Autofluorescence was reduced using Vector TrueVIEW reagent (Vector Laboratories, Newark, CA, USA; Cat# SP-8400). Positive controls were generated using DNase I (Thermo Fisher Scientific, Waltham, MA, USA; Cat# 18068-015) treatment. Micrographs were acquired using a Nikon light microscope (Model Eclipse Ni-U, 944389, Nikon Instruments Inc., Melville, NY, USA) equipped with a PCO.edge camera (Excelitas PCO GmbH, Kelheim, Germany). Quantification of TUNEL positive cells in the epithelium was performed using NIS-Elements AR software (version 6.10.02; Nikon Instruments Inc., Melville, NY, USA).

### Immunofluorescence staining

Paraffin-embedded tongue sections were deparaffinized, rehydrated, and subjected to antigen retrieval using citrate buffer (Vector Laboratories, Newark, CA, USA; Cat# H-3300-250). Sections were blocked with 4% normal donkey serum (Jackson ImmunoResearch, West Grove, PA, USA; Cat# 017-000-121) and 0.1% Triton X-100 (Sigma-Aldrich, St. Louis, MO, USA; Cat# T8787). Primary antibodies were applied overnight at 4°C, including rabbit anti-FABP3 (1:350; Thermo Fisher Scientific, Waltham, MA, USA; Cat# PA5-92386, RRID: AB_2800296), rabbit anti-CPT1C (1:350; Thermo Fisher Scientific; Cat# PA5-98783, RRID: AB_2810142), rabbit anti-BAX (1:250; Abcam, Cambridge, UK; Cat# ab32503, RRID: AB_725631), rabbit monoclonal anti-FABP5 (D1A7T) (1:350; Cell Signaling Technology, Danvers, MA, USA; Cat# 28402, RRID: AB_2799165), mouse monoclonal anti-p63 (1:50; Santa Cruz Biotechnology, Dallas, TX, USA; Cat# sc-25268, RRID: AB_628091), rabbit polyclonal anti-EpCAM (1:200; Abcam, Cambridge, UK; Cat# ab71916), mouse monoclonal anti-BCL2 (1:50; BioLegend, San Diego, CA, USA; Cat# 633502, RRID: AB_2563620), rabbit monoclonal anti-ATF3 (1:50; Abcam; Cat# ab207434, RRID: AB_2860542), mouse monoclonal anti-Waf1/Cip1/CDKN1A p21 (1:50; Santa Cruz Biotechnology; Cat# sc-6246, RRID: AB_628073), mouse monoclonal anti–c-Jun (1:50; Santa Cruz Biotechnology; Cat# sc-74543, RRID: AB_628115), and mouse monoclonal anti-Krt8 (1:50; Developmental Studies Hybridoma Bank, Iowa City, IA, USA; TROMA-I, RRID: AB_531826). Secondary antibodies included goat anti-mouse Alexa Fluor 546 (1:500; Thermo Fisher Scientific, Waltham, MA, USA; Cat# A-11003, RRID: AB_2534077) and donkey anti-rabbit Alexa Fluor Plus 488 (1:500; Thermo Fisher Scientific; Cat# A32790, RRID: AB_2762834). Nuclei were stained with Hoechst 33342 (Thermo Fisher Scientific; Cat# H3570), and slides were mounted using ProLong Gold Antifade Mountant (Thermo Fisher Scientific; Cat# P36934). Epifluorescence micrographs for FABP3, FABP5, and CPT1C were acquired using a Nikon fluorescence microscope (Model Eclipse Ni-U, 944389, Nikon Instruments Inc., Melville, NY, USA) equipped with a PCO.edge camera and controlled by NIS-Elements AR software (version 6.10.02; Nikon Instruments Inc., Melville, NY, USA). All other markers were imaged using a Dragonfly spinning disk confocal microscope (Andor Technology, Oxford Instruments, Belfast, UK). Imaging settings were kept constant across groups.

### Western blot assays

Protein extracts were prepared from tissues homogenized on ice in radioimmunoprecipitation assay (RIPA) buffer (MilliporeSigma, Burlington, MA, USA; Cat# R0278) supplemented with a protease inhibitor cocktail (Protease Inhibitor Cocktail, MilliporeSigma; Cat# P8340) and phosphatase inhibitor cocktails 1 and 2 (MilliporeSigma; Cat# P2850 and P5726, respectively) at 1:100 dilution. Homogenization was performed using Lysing Matrix D tubes (MP Biomedicals, Irvine, CA, USA; Cat# 116913050) in a FastPrep-24™ 5G instrument (MP Biomedicals, Irvine, CA, USA; Cat# 116005500) at 6.0 m/s for 40 s, with cooling on ice between cycles. Lysates were centrifuged (2795 x *g*, 1 min, 4°C), incubated on ice for 30–40 min, and further clarified by centrifugation (16099 x *g*, 40 min, 4°C). Protein concentrations were determined using the Pierce bicinchoninic acid (BCA) Protein Assay Kit (Thermo Fisher Scientific, Waltham, MA, USA; Cat# 23227) with bovine serum albumin standards (Thermo Fisher Scientific, Waltham, MA, USA; Cat# 23209; 20–2000 μg/mL) and normalized across samples. Samples were mixed with Laemmli sodium dodecyl sulfate (SDS) sample buffer (2× Laemmli Sample Buffer, Bio-Rad, Hercules, CA, USA; Cat# 1610737EDU) containing β-mercaptoethanol (Sigma-Aldrich, St. Louis, MO, USA; Cat# M6250) and denatured at 95°C for 10 min. Proteins were separated by SDS–polyacrylamide gel electrophoresis (SDS-PAGE) using mini-PROTEAN TGX precast gels (4–20% gradient; Bio-Rad, Hercules, CA, USA; Cat# 456-1094) and transferred to nitrocellulose membranes (Bio-Rad, Hercules, CA, USA; Cat# 1620167) in Tris-glycine transfer buffer (25 mM Tris, 192 mM glycine, 20% methanol) at 100 V for 1 h at 4°C. Membranes were blocked in 5% (w/v) nonfat dry milk prepared in Tris-buffered saline with Tween-20 (TBS-T), where Tris-buffered saline (TBS) consisted of 20 mM Tris-HCl and 150 mM NaCl (pH 7.6), and TBS-T contained 0.1% (v/v) Tween-20. Membranes were incubated with a Cleaved Caspase-3 (Asp175) rabbit monoclonal antibody (Cleaved Caspase-3 (Asp175) antibody, Cell Signaling Technology, Danvers, MA, USA; Cat# 9661) and a GAPDH (14C10) rabbit monoclonal antibody (GAPDH (14C10) antibody, Cell Signaling Technology, Danvers, MA, USA; Cat# 2118; RRID: AB_561053) diluted 1:1000 at 4°C overnight. After washing, membranes were incubated with a horseradish peroxidase (HRP)-conjugated anti-rabbit IgG secondary antibody (Cell Signaling Technology, Danvers, MA, USA; Cat# 7074P2, RRID: AB_2099233) diluted 1:10,000 for 1 h at room temperature. Protein bands were visualized using enhanced chemiluminescence (SuperSignal™ West Pico PLUS Chemiluminescent Substrate, Thermo Fisher Scientific, Waltham, MA, USA; Cat# 34577) and imaged using a ChemiDoc Imaging System (Bio-Rad, Hercules, CA, USA; Cat# 17001401). Positive and negative controls included Jurkat cell extracts treated or not with cytochrome c (Cell Signaling Technology; Cat# 83979 and 64514, respectively).

### Bulk RNA sequencing analysis

Tongue and small intestine tissues were dissected and immediately submerged in RNAprotect Tissue Reagent (Qiagen, Germantown, MD, USA; Cat# 76104) and stored at −80°C until processing. Samples were centrifuged (7227 x *g*, 5 min) to remove excess reagent, and tissues were sectioned into smaller fragments. Total RNA was extracted using the RNeasy Mini Kit (Qiagen, Germantown, MD, USA; Cat# 74104) according to the manufacturer’s instructions with minor modifications. Tissue fragments were lysed in 600 μl Buffer RLT supplemented with β-mercaptoethanol (10 μl per 1 mL; 2-mercaptoethanol; β-mercaptoethanol, Sigma-Aldrich, St. Louis, MO, USA; Cat# M6250) using Lysing Matrix D tubes (MP Biomedicals, Irvine, CA, USA; Cat# 116913050), followed by homogenization in a FastPrep-24 instrument (MP Biomedicals, Irvine, CA, USA) at 6.0 m/s for 40 s. Lysates were cleared by centrifugation (2795 x *g*, 1 min), mixed with an equal volume of 70% ethanol (Thermo Fisher Scientific, Waltham, MA, USA; Cat# BP28184), and applied to RNeasy spin columns. Columns were washed and subjected to on-column DNase digestion using the RNase-Free DNase Set (50) (Qiagen, Germantown, MD, USA; Cat# 79254) for 15 min at room temperature. Following DNase treatment, columns were washed with Buffer RW1 and Buffer RPE, and RNA was eluted in 30 μl RNase-free water. RNA concentration and purity were measured using a NanoDrop spectrophotometer (Thermo Fisher Scientific, Waltham, MA, USA), and RNA integrity was assessed using a Bioanalyzer 2100 (Agilent Technologies, Santa Clara, CA, USA).

Libraries were prepared using the Illumina RiboZero Stranded Total RNA library preparation kit and sequenced on an Illumina NovaSeq 6000 and demultiplexed using the Illumina bcl2fastq version 2.20.0.422 software, to generate 2×100 paired-end fastq format files. Bulk RNA-seq data were processed using the ENCODE uniform RNA-seq pipeline (v1.2.4)^24^. Briefly, reads were aligned to the mm10 mouse genome with GENCODE M21 gene annotation^25^ using STAR^26^, and gene-level expression was quantified using RSEM^27^. Differential gene expression analysis was performed in R using DESeq2. Genes with an absolute log2 fold change (FC) ≥1 and a false discovery rate (FDR)-adjusted P value ≤ 0.05 were considered differentially expressed. Functional enrichment analysis of differentially expressed genes was performed using GOseq^28^ which accounts for gene length bias. Direct p53 targets were annotated based on literature^20,29^. Upregulated and downregulated genes were analyzed separately according to the direction of change, using all genes tested in the differential expression analysis as the background gene universe. Gene Ontology (GO) and Kyoto Encyclopedia of Genes and Genomes (KEGG) pathway enrichment analyses were performed, and P values were adjusted for multiple testing using the Benjamini-Hochberg method. Categories with FDR < 0.05 were considered significant. Redundant GO terms were summarized using REVIGO^30^.

### Single-cell RNA sequencing (scRNASeq) and analysis

Adult mouse tongues from PBS-treated and 5-FU-treated animals sacrificed at days 2 (D2), 4 (D4), and 8 (D8) were harvested and immediately processed for scRNASeq. Tongues were dissected using a scalpel to isolate the dorsal mucosa. The dissected dorsal tissue was dissociated into single-cell suspension using the Whole Skin Dissociation Kit (Miltenyi Biotec, Gaithersburg, MD, USA; Cat# 130-101-540) according to the manufacturer’s instructions. Briefly, tissues were incubated in the supplied enzyme mixture and mechanically dissociated using the gentleMACS™ Dissociator (Miltenyi Biotec, Gaithersburg, MD, USA). The resulting cell suspension was filtered to remove aggregates, washed, and assessed for cell concentration and viability by trypan blue exclusion using a Luna automated cell counter (Logos Biosystems, Fairfax, VA, USA). Samples were then processed using the PIPseq^TM^ platform (Fluent Biosciences, Watertown, MA, USA), and libraries were sequenced for downstream analysis. Cell suspensions from 2 mice were pooled for sequencing. Duplicate pools per condition were sequenced separately, except for D4 samples which were sequenced in a single batch due to low cell numbers.

Raw single-cell RNA-seq data were processed using the manufacturer-recommended workflow (PIPSeeker v01.01.07, GRCm39-2022.04 reference) and downstream analyses were performed in R using Seurat v5^31^. Cells from all groups were normalized and integrated using Seurat SCTransform, generating a single Seurat object for downstream single-cell analysis in R. Dimensionality reduction, clustering using shared nearest neighbor (SNN) graph-based clustering across multiple resolutions, and visualization by uniform manifold approximation and projection (UMAP)^32^ were performed using a standard workflow. Major cell clusters were annotated after differential expression analysis performed against all remaining cells pooled together using Seurat’s FindMarkers function. Cell types were annotated using canonical marker gene expression and grouped into major cell classes, including epithelial, immune, fibroblast, endothelial, and muscle populations. Epithelial cells were further subclustered into 22 subclusters which were annotated by 1) comparing the gene expression of each cluster against non-epithelial cells; 2) comparing the gene expression of each cluster to other epithelial clusters, and 3) literature-supported lineage markers. To complement cluster-based annotation, module scores were calculated using curated marker sets. Cells were assigned to classes according to predefined score thresholds, and cells that did not meet these criteria were classified as unassigned. Cell-type distributions across timepoints were quantified from cell counts and visualized as proportional bar plots. Within the epithelial compartment, subpopulation abundances across timepoints were similarly quantified and displayed as heatmaps of per-day proportions.

Time-dependent transcriptional changes were assessed within annotated cell types and epithelial clusters by pairwise comparison of PBS with each post-treatment timepoint (D2, D4, and D8 5-FU). Genes with adjusted P value ≤ 0.05 were considered differentially expressed. Functional enrichment of time-dependent differentially expressed genes was performed using GOseq, with upregulated and downregulated genes analyzed separately and the full set of detected genes used as the background. P values were adjusted by the Benjamini-Hochberg method and GO terms and KEGG pathways with FDR < 0.05 were considered significant. Redundant GO terms were summarized using REVIGO^30^.

To evaluate co-expression within single cells of selected gene pairs leading to either apoptosis or cell cycle arrest, cells were grouped in intervals according to the expression levels of the two genes of interest, and the percentage of cells co-expressing the two genes within each interval subgroup was then calculated. Co-expression density heatmaps were generated with each heatmap tile representing cell subgroups with defined expression levels for each gene. Color intensity of each tile indicates the percentage of cells in the corresponding subset co-expressing the two genes.

### Clinical evaluation of oral mucositis in patients undergoing cancer treatment

Clinical cases were derived from a prospective, longitudinal, observational study conducted at the Roswell Park Comprehensive Cancer Center. Adult patients (≥21 years of age) were eligible for enrollment if they were scheduled to initiate a new cycle of chemotherapy, with or without concomitant radiation therapy, for solid tumors of the breast, gastrointestinal tract, head and neck, or thorax. Eligible patients were enrolled prior to initiation of their planned treatment cycle. Participants were followed longitudinally during treatment for up to 45 days. At each study visit, standardized oral examinations were performed to assess the development and progression of mucositis. The study protocol was approved by the Institutional Review Board at Roswell Park Comprehensive Cancer Center (IRB protocol #I-2983022). All procedures were conducted in accordance with institutional guidelines and applicable regulatory standards.

## Results

### Evaluation of temporal dynamics of 5-FU–induced gastrointestinal mucosal injury reveals epithelial atrophy as the main feature of the response

5-FU was administered to mice via intraperitoneal injections following a sub-lethal administration scheme for 6 consecutive days that allowed development of upper and lower gastrointestinal injury (Fig. 1A). Animals showed progressive weight loss during 5-FU administration that continued for 2 days following 5-FU withdrawal (Supplemental Fig. S1A). Oral lesions, stained with toluidine blue to indicate areas of epithelial erosion, were observed from day 4 and continued to be visible until day 10 of the follow-up period (Figs. 1B-C). Tongue tissues showed progressive loss of the apical keratinized epithelial layer, overall epithelial thinning and loss of basal cells, beginning as early as day 2 and progressing to loss of at least one third of epithelial thickness and depletion of half of basal cells by day 4, continuing until day 8 (Figs. 1D-F). These changes were followed by restoration of epithelial thickness and basal cell density by day 10, highlighting the rapid tissue recovery after removal of 5-FU (Figs. 1D-F). Histological tracking of toluidine-blue-positive tongue areas showed marked toluidine blue stained areas coincided with superficial intra-epithelial clefts or deeper breaches that exposed the submucosa (Supplemental Fig. S1B).

**Figure 1.**
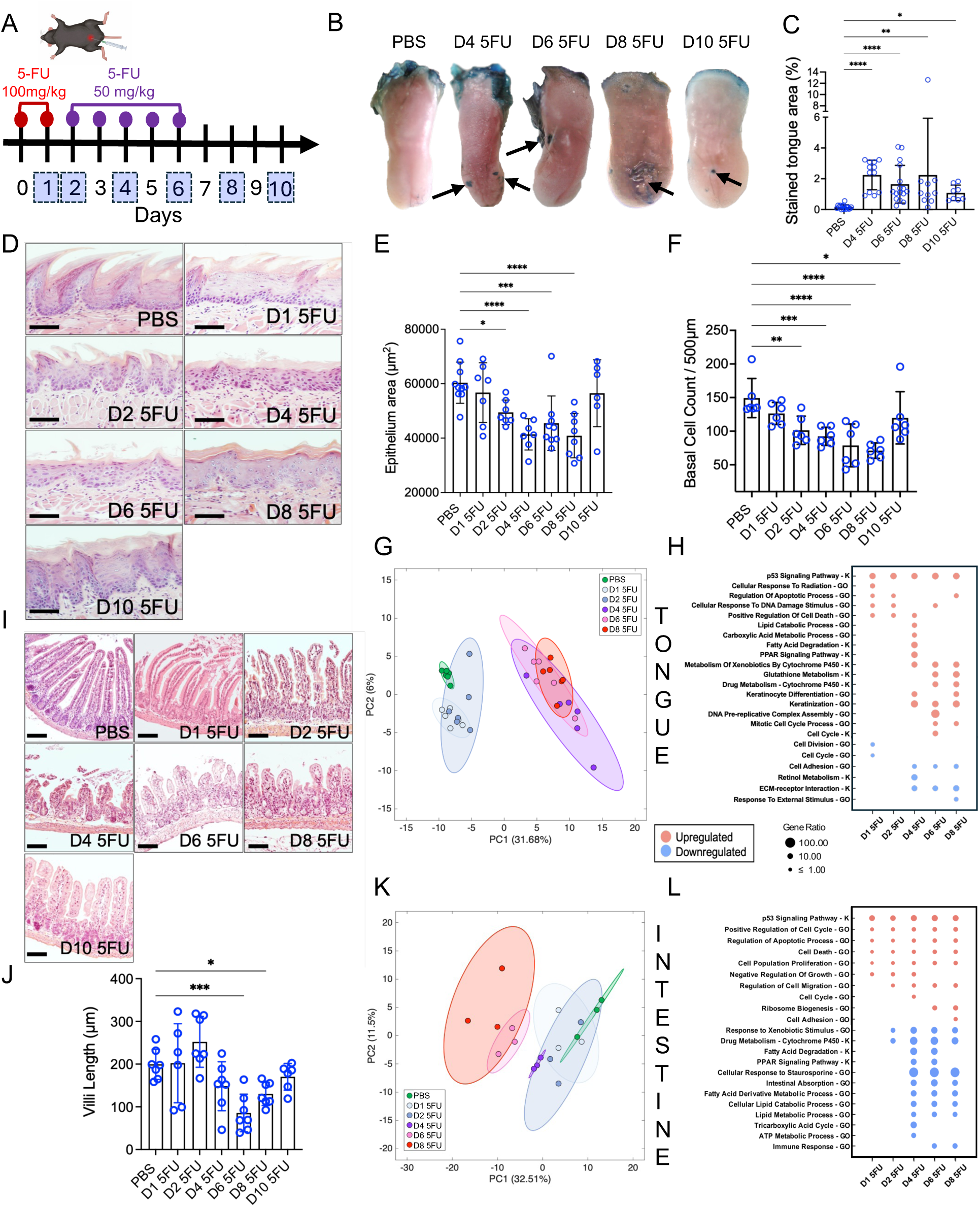
5-FU induces progressive epithelial tissue injury and transcriptional remodeling in oral and intestinal mucosae. (A) Experimental scheme of 5-FU administration with numbers in blue squares indicating mouse sacrifice days. (B) Representative images of tongues from PBS- and 5-FU-treated mice showing loss of epithelial integrity as indicated by toluidine blue staining. (C) Quantification of tongue dorsum toluidine-blue stained lesions. (D) Representative histological micrographs of tongue dorsum sections stained with hematoxylin and eosin (H&E) from PBS- and 5-FU-treated mice (Scale bar = 50 µm). n = 6 mice per group. (E) Quantification of changes in epithelial thickness in tongue dorsum in response to 5-FU. (F) Quantification of changes in basal cell counts in tongue dorsum response to 5-FU. (G) Principal component analysis (PCA) plot of tongue transcriptomes in PBS- and 5-FU-treated mice as determined via bulk RNA sequencing. n = 6 mice per group. (H) Gene Ontology [GO] term and KEGG [K] pathway enrichment analyses of tongue transcriptome evaluated via bulk RNA sequencing, comparing 5-FU-exposed mice to PBS controls. (I) Representative histological micrographs of small intestine sections stained with H&E (Scale bar = 50 µm). n = 6 mice per group. (J) Quantification of intestinal villi length in PBS- and 5-FU-treated mice. (K) Principal component analysis (PCA) plot of small intestinal tissue transcriptome in PBS- and 5-FU-treated mice as analyzed via bulk RNA sequencing. n = 3 mice per group. (L) Gene Ontology [GO] term and KEGG pathway [K] enrichment analyses of small intestine comparing 5-FU-exposed mice to PBS controls. Data in C, E, F and J are presented as mean ± SD; *p<0.05, **p<0.01, ***p<0.001, ****p<0.0001, as determined via ANOVA with Tukey post hoc analysis.

The loss of the tongue apical keratinized layer containing irregular papillary projections in mice (Fig. 1D) corresponds to clinical changes seen in patients undergoing chemotherapeutic treatment for cancer, in which loss of the “brush-like” appearance of the tongue dorsum, also known as the filiform papillae, is one of the first observed signs of mucositis (Supplemental Fig. S2A-C). In humans, the tongue dorsum could also develop erythema and mucosal breaches, with multiple oral sites other than the tongue often affected.

Evaluation of the mouse intestinal mucosa mirrored the histological changes seen in the oral cavity, exhibiting significant shortening of villi and crypt collapse during the injury phase, with partial structural restitution after 5-FU withdrawal (Fig. 1I-J). In both tongue and intestine, the connective and muscular layers were preserved, suggesting that 5-FU injury is highly compartment-specific and targets the epithelium rather than deeper tissue strata.

Histological staining of oral and intestinal tissues did not suggest accumulation of an inflammatory infiltrate in response to 5-FU. To confirm this, we performed immunohistochemical staining for the myeloid cell surface marker CD45. We observed a decrease in CD45-positive cells in the oral mucosa during injury and no changes in the number of CD45-positive cells in the intestine apart from a decrease at day 10 (Supplemental Fig. S3A-D). Altogether, these results suggest that the main tissue compartment affected by 5-FU is the epithelium, which experiences cell loss, atrophy and structural damage.

### Oral and intestinal tissues show common p53 signaling induction and site-specific transcriptional responses to 5-FU

Next, we performed bulk RNA sequencing to examine the nature of responses to 5-FU in tongue and intestinal mucosal tissues. This analysis revealed a temporal trajectory of gene expression that paralleled the observed morphological changes (Supplemental Tables S1 and S2). Principal component analysis (PCA) of oral samples showed 5-FU-treated animals examined at days 1 and 2 clustered together with PBS-treated controls, with a marked shift in global gene expression occurring after day 4 (Fig. 1G). The early tongue response at days 1 and 2 was characterized by upregulation of the p-53 signaling pathway, and related GO terms such as “positive regulation of cell death”, “regulation of apoptotic process” and “cellular response to DNA damage stimulus”, accompanied by a downregulation of cell division and cell cycle (Fig. 1H). Activation of p53 signaling continued throughout all the days evaluated. However, at day 6 we also saw evidence of a reactivation of the cell cycle suggesting a recovery in cell proliferation. In addition, we observed upregulation of lipid, more specifically fatty acid, degradation at day 4 suggestive of a switch in cellular metabolism.

Similarly to tongue tissues, the intestine showed a temporal shift in global gene expression (Fig. 1K) and upregulation of p53 signaling on all experimental days (Fig. 1L). However, in contrast to the tongue, intestinal tissues showed from day 4 a downregulation in metabolic processes involved in energy production and lipid catabolism. Altogether, these results suggest the transcriptional regulator p53 orchestrates the response to 5-FU in both oral and intestinal compartments, but tissues also display site-specific responses related to cellular metabolism.

### 5-FU induced p53-dependent but compartment-specific gene signatures suggestive of cell cycle arrest and apoptosis as mediators of tissue damage

The p53 transcriptional regulator is known for determining cell fate decisions in response to cellular stress via activation of context-specific target genes^33–35^. p53 can induce DNA damage repair, cell cycle arrest and/or apoptosis, as well as regulate other processes including energy metabolism. However, the way p53 controls cell fate, especially in vivo, in different tissues and cell types is incompletely understood. To better delineate the type of transcriptional p53-dependent program induced by 5-FU in oral and intestinal mucosae, we identified direct p53 transcriptional targets differentially regulated in our bulk RNA-seq datasets and grouped them based on their function (Fig. 2A). This analysis showed that genes related to cell cycle and apoptosis were both prominent responses induced in tongue and intestine, however, each tissue showed a specific signature of regulated genes. For instance, while both tongue and intestine showed upregulation of the growth suppressor *Psrc1* at all time points, the master cell cycle regulator, *Cdkn1a* (*p21*) - which inhibits cell cycle progression by binding to different cyclin/Cdk1 complexes^36^- was upregulated throughout the time course in intestine, but was only significantly upregulated at days 1 and 2 in tongue. In tongue, coinciding with the restricted upregulation of *Cdkn1a*, related cell cycle genes including the cyclin B complex *Ccnb1* and *Ccnb2*, the master cell cycle regulator *Cdk1* and the *Cdkn1a* regulator *Cdc20* were all downregulated at days 1 and 2 and then upregulated at day 4. These results suggest that the tongue mucosa responded to the 5-FU challenge by temporarily inducing a transcriptional program compatible with cell cycle arrest, followed by later transcriptional responses supportive of cell cycle re-entry and replication. These responses were less clearly delineated in the intestine, where *Cdkn1a* (*p21*) showed increased expression at all time points but the concerted downregulation of genes that promote progression through the cell cycle was not observed. Instead, the less studied growth suppressor gene *Serpinb5* was induced during all time points in intestine where it could be playing a tissue-specific role.

**Figure 2.**
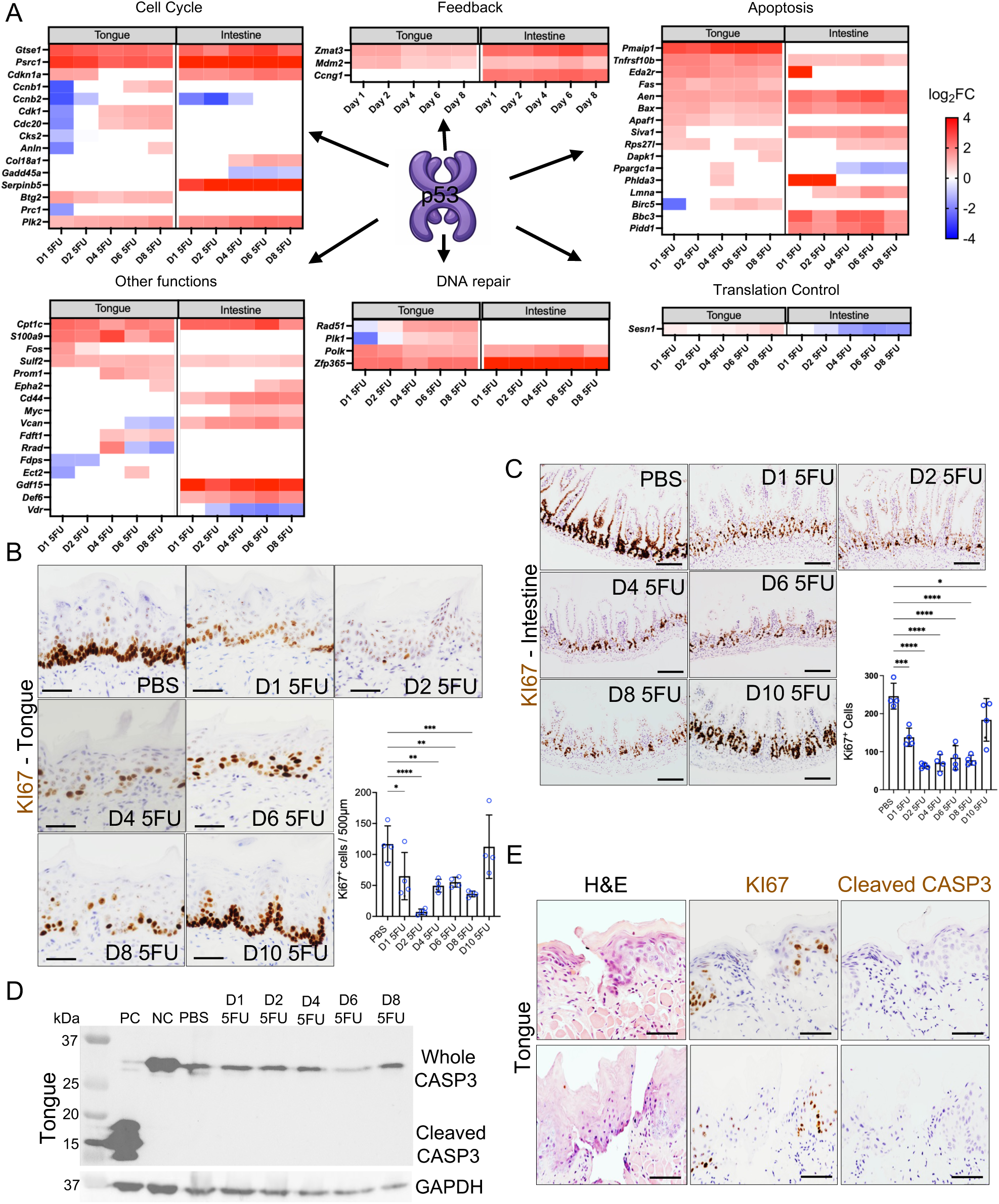
Common and compartment-specific p53-driven transcriptional signatures include genes involved in controlling divergent cell fates with cell cycle arrest as the dominant tissue response. (A) Heatmaps showing differentially expressed p53 targets in tongue and intestine in 5-FU-treated mice compared to PBS as determined via bulk RNA sequencing. Only genes with log2 fold-change (log2FC) > 1 or < −1 and adjusted p values < 0.05 are shown. (B-C) Representative micrographs and quantification of KI67 staining indicating proliferative cells in tongue mucosa (B) and small intestine (C) of PBS- and 5-FU-treated mice. (D) Western blot of total and cleaved Caspase-3 protein in tongue tissue lysates of PBS- and 5-FU-treated mice. Positive control (PC) is Jurkat cells treated with 1 μM staurosporine, and Negative control (NC) is untreated Jurkat cells. (E) Representative micrographs of sections corresponding to tongue lesion areas in mice treated with 5-FU and sacrificed at day 4 showing H&E staining and immunohistochemistry for detection of KI67 and cleaved Caspase 3 protein (Scale bar = 50 µm). Data in B and C are presented as mean±SD; *p<0.05, **p<0.01, ***p<0.001, ****p<0.0001, as determined via ANOVA with Tukey post hoc analysis.

The analysis of p53 targets also showed a strong pro-apoptotic transcriptional response to 5-FU in tongue and intestine (Fig. 2A). Critical intrinsic apoptosis effectors, such as *Bax* and *Aen* and extrinsic apoptosis activators including *Tnfrsf10b* (Death Receptor 5) were upregulated in both tissues throughout the time course. However, apoptosis-related compartment-specific responses were also evident with upregulation of pro-apoptotic genes such as *Pmaip-1* (*Noxa*), *Apaf-1* and *Fas* only in tongue, while *Bbc3* (*Puma*) and *Pidd1* were intestine-specific.

### Cell cycle arrest is the dominant response to 5-FU in tongue and intestine

Since it appeared that 5-FU simultaneously induced transcriptional programs leading to competing cell fates, we next evaluated cell proliferation and apoptosis to determine the presence and temporal dynamics of these processes in tongue and intestinal mucosae. Immunohistochemical staining for the proliferation marker KI67 in tongue tissues revealed a sharp decline in actively cycling epithelial basal cells from day 1, with almost no KI67-positive cells remaining at day 2 (Fig 2B). By day 4, KI67 expression started to recover, achieving normal values at day 10 (Fig 2B). These changes were similar in intestinal crypts, where 5-FU treatment suppressed KI67 positivity in the proliferative zone (Fig. 2C). However, contrary to the complete obliteration of KI67 expression observed in tongue by day 2, a small number of cells expressing high levels of KI67 remained in the intestine throughout 5-FU treatment. Moreover, although on their way to recovery, intestinal tissues could not reach normal proliferation levels by day 10, suggesting the oral epithelium was more resilient to the effects of 5-FU than the small intestine.

In contrast, immunohistochemical staining for cleaved Caspase-3 to detect apoptosis revealed only scattered single positive cells in both tongue and intestines with no statistically significant increase in staining during 5-FU treatment (Supplemental Figs. S4A-D). We validated these results by performing TUNEL assay on tongue tissues to detect DNA fragmentation due to apoptosis. This assay also showed low levels of DNA fragmentation and no increase in TUNEL-positive cells after 5-FU treatment (Supplemental Fig S4E-F). Western blot assays confirmed stable levels of full-length Caspase-3 and absence of the 17 kDa cleaved fragment in tongue tissues after 5-FU treatment (Fig. 2D). Fig. 2E shows that tongue areas with epithelial breaches show a depletion of KI67-positive proliferative cells around the lesion without evidence of cleaved Casapase-3-positive apoptotic cells. Collectively, these results show that although 5-FU induced a p53-driven transcriptional response involving regulators of cell cycle arrest and apoptosis, the dominant response leading to tissue damage is the former.

### Single-cell transcriptomics of tongue confirms the epithelial compartment as the principal target of 5-FU and identifies epithelial subpopulations with different susceptibilities to the drug

Next, we focused our analysis on the oral mucosa, a tissue in which the progenitor populations and tissue responses to injury are largely uncharacterized, by subjecting dissected tongue dorsum mucosal tissue to scRNASeq. For these experiments, 5-FU was withdrawn after day 4 to allow evaluation of recovery populations by day 8 (Fig. 3A). scRNASeq identified 5 distinct cell populations in uninjured mucosa corresponding to broad cell types (epithelial, fibroblasts, endothelial, muscle and immune) (Fig. 3B, Supplemental Table S3 and Supplemental Fig S5A). Proportions of the cell types in the sequenced dataset changed after 5-FU exposure (Fig. 3C and Supplemental Fig. S5B). Consistent with previous histological findings, epithelial cells were depleted through days 2 and 4 but their proportion fully recovered after 5-FU removal.

**Figure 3.**
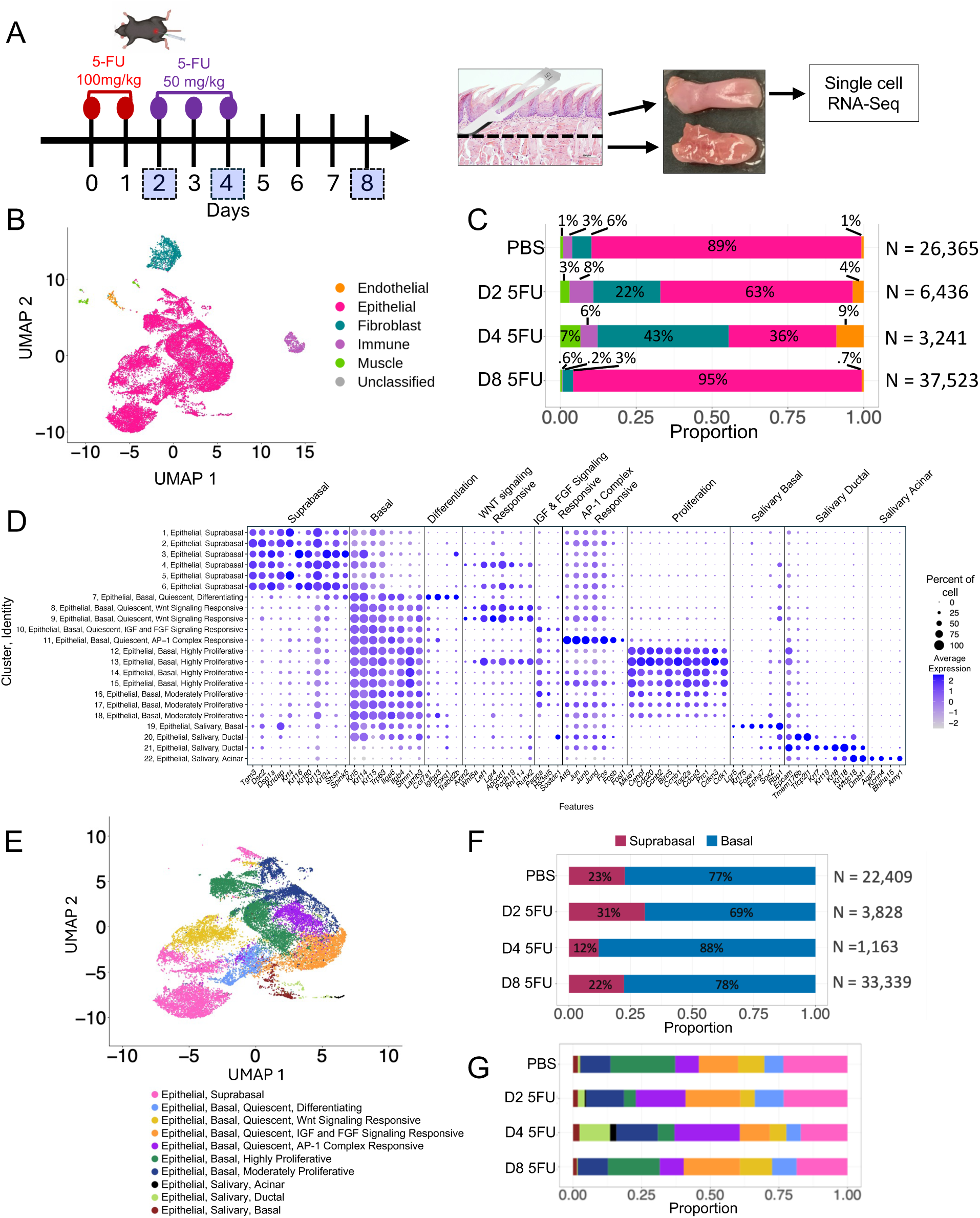
Single cell (sc)RNASeq of tongue tissues identifies epithelial subpopulations with different susceptibility to 5-FU. (A) Experimental scheme showing schedule followed for intraperitoneal 5-FU administration with numbers in squares indicating mouse sacrifice days. For scRNASeq the tongue dorsal surface was dissected. (B) Uniform Manifold Approximation and Projection (*UMAP*) showing major cell populations identified in tongue dorsum. (C) Longitudinal cell composition analysis showing changes in proportion of major cell types in PBS and 5-FU treated mice across experimental time points. (D) Dot plot depicting expression of main gene markers in epithelial subclusters, indicating suprabasal, basal and different epithelial cell subpopulation identities. Bubble size represents percentage of cells expressing the gene and color intensity represents average gene expression. (E) UMAP showing epithelial subpopulations identified. (F) Longitudinal changes in proportion in epithelial subpopulations after 5-FU administration.

We next focused on epithelial cells to identify subpopulations and evaluate their dynamics in response to 5-FU. We identified 22 subclusters within the epithelial compartment (Supplemental Fig. S6A). Each epithelial subcluster was compared to non-epithelial and to other epithelial cells to establish identifying markers, which were validated by literature searches (Supplemental Fig. S6B, Supplemental Table S4). Based on this analysis, epithelial cells could be divided into three major subtypes with suprabasal, basal and salivary gland identities (Figs. 3D-E).

An evaluation of changes in the proportions of suprabasal and basal epithelial cells after 5-FU exposure showed depletion of basal populations at day 2, while at day 4 cells with a suprabasal identity decreased, with both populations re-establishing their original ratio by day 8 (Figure 3F and Supplemental Fig. S7). These dynamics suggest that basal cells were highly susceptible to the early high dose of 5-FU and highlight the ability of the oral epithelium to rapidly recover from chemotoxic injury.

Next, we assigned identities based on gene signatures to the different subpopulations of epithelial basal cells and evaluated how their proportions changed after 5-FU administration. Based on proliferation markers, we identified quiescent (clusters 7-11), highly proliferative (Clusters 12-15) and moderately proliferative (Clusters 16-18) cells (Fig. 3D). Among quiescent basal cells, there were different subpopulations identified based on their gene signatures and labelled as differentiating (Cluster 7), Wnt-signaling responsive (Clusters 8 and 9), Insulin Growth Factor (IGF) and Fibroblast Growth Factor (FGF) signaling responsive (Cluster 10), and a population of Activator Protein-1 (AP-1) complex responsive cells (Cluster 11) (Figs. 3D, 3E and Supplemental Table S4). The Wnt-signaling responsive cells appear as plausible epithelial progenitor candidates as Wnt is a critical pathway in progenitor populations that control homeostasis in other epithelial tissues^37,38^. Furthermore, a cluster of basal proliferative cells (Cluster 13) and clusters of suprabasal cells (Clusters 4 and 6) also showed a preserved Wnt gene signature (Fig. 3D), representing potential progeny.

Evaluation of changes in the proportions of basal subclusters showed depletion of the highly proliferative population at days 2 and 4, while moderately proliferative cells were more resistant to 5-FU (Fig. 3G and Supplemental Figs. S8A-B). Among quiescent cells, the Wnt-signaling-responsive cells showed early depletion at day 2, while other quiescent populations appeared resistant at this early time point. At day 4, all quiescent populations were depleted compared to PBS control tissues except the AP-1 complex responsive population which became the dominant epithelial population (Fig. 3G).

Epithelial cells also included salivary basal (Cluster 19), salivary ductal (Clusters 20, 21) and salivary acinar (cluster 22) cells identified based on their gene signatures (Figs. 3D and E). Minor salivary glands (Von Ebner’s glands) are known to exist in the tongue embedded within the connective tissue^39^. Epcam which has widespread expression in simple, glandular and pseudostratified epithelium but is absent in stratified epithelia such as skin^40^, was one of the markers for these salivary gland cells. To confirm the presence of salivary glands in tongue, we performed EPCAM immunostaining. As seen in Supplemental Fig. S9, strong EPCAM staining revealed salivary gland structures in the tongue while it exhibited faint staining in other tongue epithelial cells. Salivary gland clusters appeared resistant to the effects of 5-FU (Fig. 3G and Supplemental Figs. S8A-B). Of note, we did not observe in this dataset cells with markers indicating a test bud identity^41^.

### Single tongue epithelial cells upregulate pro-apoptotic, anti-apoptotic and cell cycle arrest effectors in response to 5-FU

To further understand the responses of individual cellular compartments and cell subpopulations to 5-FU, we performed gene expression analysis based on the scRNASeq data. Across all cell types in tongue, the most extensive transcriptional remodeling occurred within epithelial cells (Figure 4A), which exhibited strong engagement of the p53 signaling pathway, early downregulation and later upregulation of cell-cycle and upregulation of lipid metabolism (Figure 4B), mirroring gene expression changes seen by bulk RNA-seq of whole tongues (Fig. 1H). Assessment of differential regulation of direct p53 targets, showed this transcriptional activation mainly occurred in the epithelium, which regulated genes related to apoptosis, cell cycle and metabolic processes (Fig. 4C and Supplemental Figs. S10A-D).

**Figure 4.**
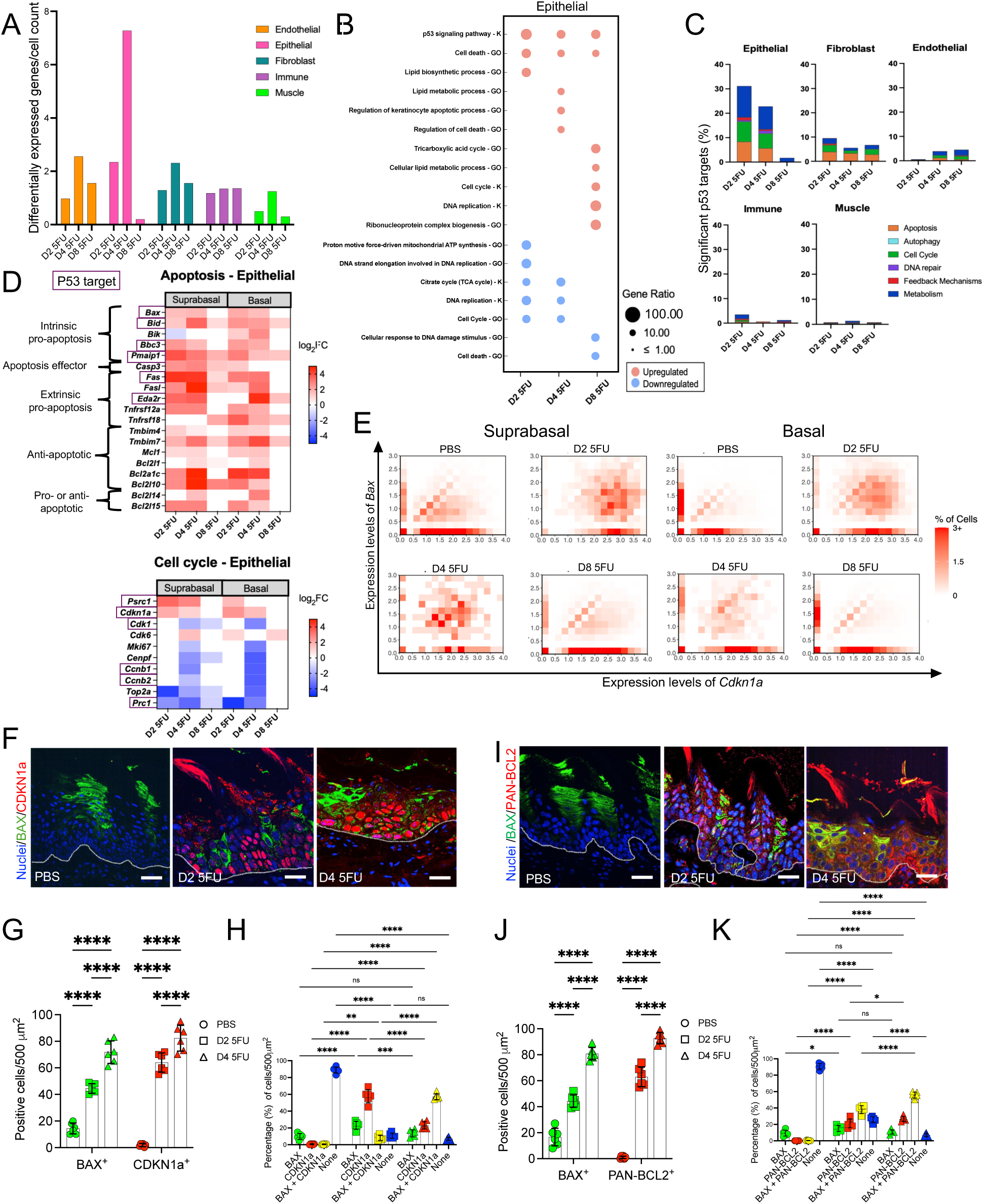
Pro-apoptotic, anti-apoptotic and anti-proliferative transcriptional programs are co-activated within single epithelial cells upon 5-FU administration. (A) Number of differentially regulated genes, according to scRNASeq, in 5-FU treated mice compared to PBS across different cell types. (B) Gene Ontology [GO] term and KEGG pathway [K] enrichment analyses in epithelial cells after 5-FU exposure in comparison to PBS, according to scRNASeq. (C) Percentage of p53 targets differentially expressed in different cell types as identified via scRNASeq. (D) Heatmaps depicting differentially expressed genes involved in apoptosis or cell cycle control in basal and suprabasal epithelial cells as evaluated via scRNASeq. Only genes with log2 fold-change (log2FC) > 1 or < −1 and adjusted p values < 0.05 are shown. Purple outline indicates p53 target. (E). Cell-level co-expression density heatmaps of the p53 target genes *Bax* and *Cdkn1a* in epithelial basal and suprabasal compartments under each condition. Cells were grouped according to *Bax* and *Cdkn1a* expression levels, and each heatmap tile represents a defined co-expression range. Color intensity indicates the percentage of cells in the corresponding compartment within each range (F-H) Representative micrographs and quantification of immunofluorescence co-staining of tongue tissues for BAX (green) and CDKN1A (red) across PBS, D2 5FU and D4 5FU conditions. Nuclei were stained with Hoechst 33342 (blue). (Scale bar = 20 µm). White line = division between epithelium and connective tissue. n = 3 mice per group. (I-K) Representative micrographs and quantification of immunofluorescence co-staining of tongue tissues for BAX (green) and pan-BCL2 (red) across PBS, D2 5FU and D4 5FU conditions. Nuclei were stained with Hoechst 33342 (blue). (Scale bar = 20 µm). White line = division between epithelium and connective tissue. N= 3 mice per group. Data in G, H, J and K are presented as mean ± SD; *p<0.05, **p<0.01, ***p<0.001, ****p<0.0001, as determined via ANOVA with Tukey post hoc analysis.

Cell cycle arrest and apoptosis are cell fates requiring different transcriptional programs. To explore the possibility that these responses are discretely upregulated in epithelial cell subpopulations at unique stages of differentiation, we compared regulation of apoptosis and cell cycle-related genes in suprabasal and basal cells, irrespective of their direct dependency on p53. As Fig. 4D shows, both basal and suprabasal cells showed parallel regulation of genes in both apoptosis and cell cycle functional categories.

To explore whether single cells simultaneously regulate genes related to divergent cell fates, we performed a within cell co-expression analysis of pairs of genes regulating either cell cycle arrest or apoptosis. To evaluate cell cycle arrest, we used the master regulator *Cdkn1a* and compared its co-expression with different pro-apoptotic genes. When co-expression of *Cdkn1a* and the critical executor of intrinsic apoptosis *Bax* was assessed, we observed an increase in cells co-expressing them within both suprabasal and basal compartments upon 5-FU administration (Fig. 4E). We validated these results at the protein level by performing immunofluorescence staining which showed BAX expression in PBS-treated tissues and an increase in levels upon 5-FU treatment, while CDKN1A was barely detectable in control tissues and its levels increased dramatically after 5-FU administration (Fig. 4F-G). Co-expression of both CDKN1A and BAX proteins in single cells was confirmed to occur with co-expressing cells seen to significantly increase after 5-FU exposure (Fig. 4H). Similar transcriptional co-expression patterns within cells were observed for *Cdkn1a* and 9 additional apoptosis-related genes (Suppl. Fig. 11A-I) confirming that upon 5-FU stimulation, single cells activate responses leading to cell cycle arrest and apoptosis, although phenotypically cell cycle arrest is the dominant response.

Analysis of gene expression in the suprabasal and basal epithelial cell compartments also showed upregulation of anti-apoptosis Bcl2-family effectors, including *Mcl1*, *Bcl2l1*, *Bcl2a1c* and *Bcl2l10* (Fig. 4D). Immunostaining with a pan-BCL2 antibody showed increased expression after 5-FU treatment (Fig. 4I-J) and co-expression of pan-BCL2 and BAX (Fig. 4I-K), showing that simultaneous expression of both pro-apoptotic and anti-apoptotic regulators occurs within tongue epithelial cells in response to 5-FU, which could explain the unexecuted apoptosis program (Fig. 4I-K).

Altogether, these results show that although a pro-apoptotic transcriptional program is activated in tongue epithelial cells during 5-FU-induced stress, simultaneous upregulation within cells of anti-apoptotic effectors and a strong upregulation of the cell cycle arrest inducer *Cdkn1a* (*p21*) result in interruption of the proliferative capacity of the epithelium rather than in cell death.

### An AP-1 complex–responsive basal epithelial subcluster with a stress-related gene signature replaces most epithelial cells by day 4 of 5-FU exposure

We next explored the basal epithelial subpopulation of AP-1 complex responsive cells, which showed the highest resistance to 5-FU. Markers for this cell population include the transcription factors *Jun*, *Junb*, *Jund*, *Fos*, *Fosb*, *Fosl1*, and *Atf3* (the AP-1 complex) (Fig. 3D), which are involved in stress responses, cell fate and reprogramming to pluripotency^42,43^. Furthermore, some of these marker genes are also direct targets of p53^29^. Additional genes enriched in the AP-1 complex responsive cell population when compared to the rest of basal cells included other p53 targets involved in adaptive stress regulation, including *Btg2*, *Ier2*, *Ier3*, *Ier5*, *Dusp1* and *Gadd45a*^44–47^ (Fig. 5A). Epithelial progenitors with a p53 and AP-1 complex gene signature have been identified in other tissues such as the lung with these pathways shown to be implicated in the maintenance of transitional cell states that enable regeneration^23^. These findings suggest this population of AP-1 complex-responsive cells may represent oral mucosal progenitors with yet uncharacterized roles in responses to injury.

**Figure 5.**
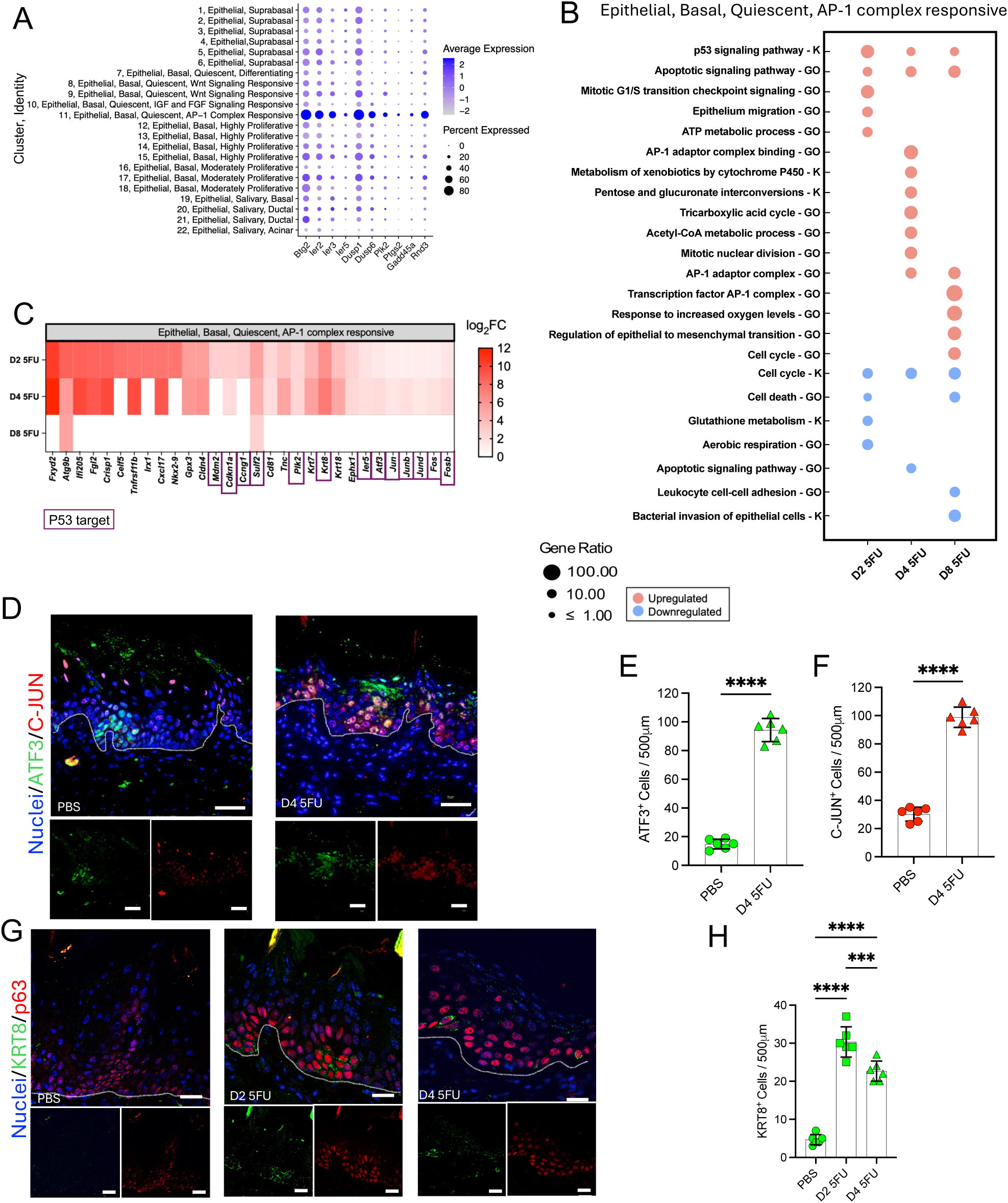
A putative oral epithelial cell progenitor population with a stress-adaptation p53 and AP1 complex-responsive gene signature persists following 5-FU exposure. (A) Bubble plot showing p53 targets enriched in the AP-1 responsive basal epithelial subpopulation during homeostasis (PBS-treated mice). Bubble size represents percentage of cells expressing gene and color intensity represents average gene expression. (B) Gene Ontology [GO] term and KEGG [K] pathways enriched in the epithelial basal AP-1–responsive subpopulation in response to 5-FU as determined via scRNASeq. (C) Heatmap showing the most differentially upregulated genes upon 5-FU exposure in the AP-1–responsive subpopulation. Only genes with log2 fold-change (log2FC) > 1 or < −1 and adjusted p values < 0.05 are shown. Purple outline indicates p53 target. (D) Representative micrographs of immunofluorescence staining of tongue tissues for the AP1-complex proteins ATF3 (green) and C-JUN (red) across PBS, D2 5FU and D4 5FU conditions (n = 3 mice per group). Nuclei were stained with Hoechst 33342 (blue). (Bar = 50µm). White line = division between epithelium and connective tissue. (E) Quantification of ATF3-positive epithelial cells. (F) Quantification of C-JUN-positive epithelial cells. (G) Representative micrographs of immunofluorescence staining of tongue tissues for the stress-response protein KRT8 (green) and the basal cell marker P63 (red) across PBS, D2 5FU and D4 5FU conditions (n = 3 mice per group). Nuclei were stained with Hoechst 33342 (blue). (Scale bar = 50 µm). White line = division between epithelium and connective tissue. (H) Quantification of KRT8-positive epithelial cells. Data in E, F and H are presented as mean ± SD; *p<0.05, **p<0.01, ***p<0.001, ****p<0.0001, as determined via ANOVA with Tukey post hoc analysis.

We next evaluated the behavior of AP-1 complex-responsive cells during the 5-FU challenge. We observed further transcriptional upregulation of the p53 signaling pathway within this population (Fig. 5B), with enrichment of p53 target genes during 5-FU administration (Fig. 5C), including *Cdkn1a* and *Ccng1* involved in cell cycle arrest^48^, but also *Mdm2* which together with *Ccng1* functions as a negative feedback regulator of p53^49^, a suggestion that temporal control of p53 levels is important for the survival of this pool of progenitors and to their potential contribution to tissue regeneration. At day 4, AP-1 complex-responsive cells also upregulated energy-related pathways and mitosis, consistent with their ability to survive 5-FU injury (Fig. 5B). Moreover, AP-1 complex genes were further enriched upon 5-FU treatment (Fig. 5B, C), together with other genes involved in adaptive cellular stress responses, such as *Krt8*, a p53-controlled, stress-inducible, intermediate filament protein previously shown to be a marker of lung epithelium cell states in development and regeneration^21–23^ and to mediate chemoresistance in cancer cells^50^. The genes *Krt18* and *Cldn4*, also markers for lung epithelium transitional cells^23^, were also enriched after 5-FU treatment in AP-1 complex-responsive cells. Moreover, *Tnc* (Tenascin-C), an extracellular matrix protein involved in wound healing, cell adhesion and migration^51^ was also upregulated.

Immunofluorescence staining of the AP-1 complex-responsive markers C-JUN and ATF3 showed scattered staining mostly in the basal layer in control tissues but by day 4 C-JUN- and ATF3-positive cells comprised the majority of cells in the epithelium (Fig. 5D-F). KRT8 protein expression was also seen to increase after 5-FU treatment but staining was confined to a smaller number of cells than C-JUN and ATF3 (Fig. 5G-H). Together, these data suggest that in addition to the AP1-complex, p53 and stress adaptation may also play a role in the maintenance of a population of putative oral epithelial cell progenitors with a program resembling transitional cells and capable of surviving chemotoxic injury.

### Metabolic reprogramming supports epithelial proliferative recovery in oral but not in intestinal mucosa

Following the resolution of cell cycle arrest, transcriptomic and histological data suggested the oral epithelium undergoes a pronounced metabolic shift that coincides with proliferative re-entry. After 5-FU, we observed in the oral, but not in the intestinal mucosae, an upregulation of lipid metabolism, which encompasses fatty acid transportation, oxidation, and catabolic processes of other lipids (Figs. 1H, 1L). In this context, we decided to compare regulation of genes involved in lipid metabolic processes in oral and intestinal tissues. As seen in Fig. 6A, in the tongue there was upregulation of genes involved in fatty acid transport and degradation/oxidation, and in lipid biosynthesis and metabolism, while these responses did not occur in the same concerted manner in the intestine. Through scRNASeq data of the oral tissue, we observed that lipid metabolism upregulation was a response almost exclusive to epithelial cells (Fig. 6B). Single cell data showed transcriptional upregulation of lipid metabolism genes in both the suprabasal and basal epithelial compartments (Fig. 6C). Very few of the genes involved in lipid metabolism are p53 targets, suggesting this response was for the most part independent of this regulator. One exception is the p53 target gene *Cpt1c* encoding the protein carnitine palmitoyltransferase 1 C, which regulates lipid metabolic reprograming and cell adaptation to environmental stressors in cancer cells^52^. Immunofluorescence staining of tongue tissues showed CPT1C protein levels were negligible in homeostasis, while there was a marked increase in expression, especially in suprabasal cells, after 5-FU treatment (Fig. 6D-E). Tongue epithelial cells also displayed marked overexpression of the fatty acid transporters FABP3 and FABP5 after 5-FU treatment compared to control tissues, with FABP3 showing increased expression in basal cells while FABP5 was preferentially expressed in the suprabasal compartment (Figs. 6F-I). Integration of transcriptomic and protein staining data suggests that 5-FU-injured oral epithelia engage a lipid oxidative metabolic program that fuels regeneration. Increased fatty-acid import and activation of the PPAR (Peroxisome Proliferator-Activated Receptor) signaling pathway may provide ATP and biosynthetic precursors necessary for cell-cycle re-entry, mitochondrial biogenesis, and membrane synthesis during tissue rebuilding. The intestinal epithelium, however, lacked this coordinated lipid metabolic response, instead relying primarily on glycolytic and amino-acid-driven metabolism during recovery. Collectively, these findings define metabolic reprogramming as the final phase in oral epithelial adaptation to chemotoxic injury.

**Figure 6.**
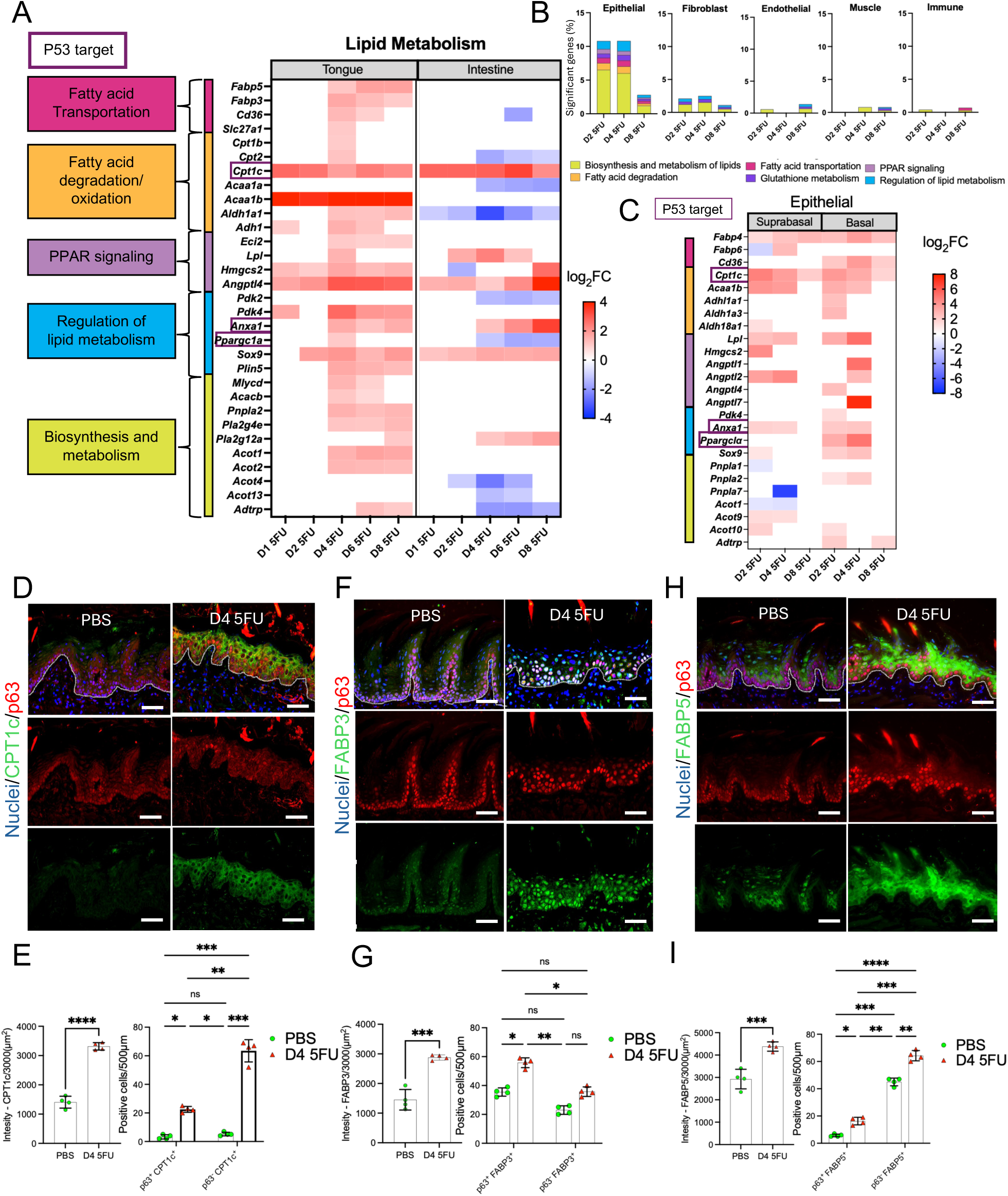
Metabolic reprogramming toward fatty-acid utilization occurs in tongue but not in intestinal tissue after 5-FU treatment. (A) Heatmap of differentially expressed genes involved in lipid metabolism in tongue and intestine as evaluated via bulk RNA sequencing. Only genes with log2 fold-change (log2FC) > 1 or < −1 and adjusted p values < 0.05 are shown. Purple outline indicates p53 target. (B) Proportion of lipid and glutathione metabolism genes significantly regulated in different cell types upon 5-FU treatment. (C) Genes involved in lipid metabolism that are differentially regulated in tongue basal and suprabasal epithelial cells as evaluated via scRNASeq. Only genes with log2 fold-change (log2FC) > 1 or < −1 and adjusted p values < 0.05 are shown. Purple outline indicates p53 target. (D) Representative micrographs of immunofluorescence co-staining of tongue tissues (n=4 mice per group) for Carnitine palmitoyltransferase 1C - CPT1C (green) and the basal marker P63 (red), in PBS and D4 5-FU conditions. Nuclei were stained with Hoechst 33342 (blue). (Scale bar = 50 µm). White line = division between epithelium and connective tissue. (E) Bar graph showing intensity of CPT1C immunofluorescent staining, and co-staining with P63 in epithelial cells. (F) Representative micrographs of immunofluorescence co-staining of tongue tissues (n= 4 mice per group) for fatty acid binding protein 3 - FABP3 (green) and the basal marker P63 (red), in PBS and D4 5-FU conditions. (G) Bar graph showing intensity of FABP3 immunofluorescent staining, and bar graph showing co-staining with P63 in epithelium. (H) Representative immunofluorescent micrographs of tongue tissue co-staining (n = 4 mice per group) for fatty acid binding protein 5 – FABP5 (green) and the basal marker P63 (red), in PBS and D4 5FU conditions. (I) Bar graphs showing intensity of FABP5 immunofluorescent staining, and co-localization with P63 in epithelium. Data in E, G and I are presented as mean ± SD; *p<0.05, **p<0.01, ***p<0.001, ****p<0.0001, as evaluated via ANOVA with Tukey post hoc analysis.

## Discussion

Mucositis due to chemotherapeutic treatment for cancer affects up to 80 % of patients undergoing cancer treatment with limited preventive or interventional therapies available^3,53,54^. 5-FU is a frequently prescribed chemotherapeutic for the treatment of various types of cancers, including gastrointestinal and head and neck carcinomas and is associated with increased mucositis severity^3,55^. Research on the pathophysiology of chemotherapy-associated mucositis is limited by the difficulty of studying the condition in humans. Although studies are available on the response of intestinal tissues to radiation-induced mucositis, there is a paucity of work addressing in vivo responses to chemotherapeutics, and there is even a wider knowledge gap when considering the response in oral tissues. Here, we employed a mouse 5-FU-based model of chemotoxic injury that replicates features of mucositis documented in humans including oral lesions, loss of tongue filiform papillae and oral and small intestinal epithelial atrophy^8,11^. This model was modified from previously published work^9^, by optimizing the 5-FU dosage to consistently observe oral lesions after intraperitoneal 5-FU administration while maximizing animal viability. We performed a broad phenotypic and transcriptional characterization of the temporal responses in the oral and small intestinal mucosae to 5-FU to gain a better understanding of common and site-specific reactions to chemotoxic stress and then focused on oral tissues performing a single cell transcriptomic evaluation. Our work provides a comprehensive map of the gene regulatory networks activated in vivo in non-tumor tissues in response to 5-FU and uncovers the landscape of cell populations in the oral mucosa outlining their kinetic responses to the drug.

Results from this study show cell cycle arrest, and not apoptosis is the key response to 5-FU in oral and intestinal tissues. Our study agrees with previous work showing decreased epithelial proliferation as evidenced by lower expression of KI67 in both tongue and intestine upon 5-FU exposure^9^. An earlier study in mice showed that apoptosis affects a very small number of cells in the intestinal mucosae after 5-FU treatment, with the number of apoptotic cells peaking one day after exposure to the drug^10^, while a study in humans detected an early spike in apoptotic cells in the small intestine one day after chemotherapy administration^11^. Agreeing with these observations, here we detected a small number of apoptotic cells in the small intestine with a trend for an increase after one day of 5-FU exposure but the difference to the PBS control did not reach statistical significance. Previous work in models of intestinal mucositis due to targeted radiation has shown, however, that deletion of critical apoptosis effectors (*Bak1* and *Bax*) from gastrointestinal epithelial cells decreases apoptosis in the small intestine but does not increase mouse survival^17^, suggesting apoptosis is not a key event that mediates small intestine damage in response to radiation. Although similar studies do not exist using chemotherapeutic drugs as the DNA damaging stimulus, the lack of a robust apoptosis response to 5-FU in the intestine suggests this cell death event is also not critical in chemotherapy-induced intestinal mucositis. Furthermore, apoptosis was even more sparsely detected in oral tissues, where we observed no increase after 5-FU treatment, although we utilized three different techniques and appropriate controls to detect it. Apoptosis, therefore, does not appear to be a mediator of tissue damage in 5-FU-induced oral mucositis.

Transcriptional analysis of the responses to 5-FU in oral and intestinal tissues showed a p53-coordinated response in both sites. Mechanistic studies using gene deletion mouse models have shown that cell cycle arrest in the intestinal mucosa after 5-FU exposure is dependent on p53^10^. Studies using sub-total body radiation targeting the intestine also show the response in this compartment is dependent on the expression of p53 and its transcriptional target *Cdkn1a*, p21, a cyclin dependent kinase-inhibitor that induces growth suppression, and although these regulators induce cell cycle checkpoints that lead to loss of tissue structure, the net effect is protective allowing recovery from injury^17,19^. Here we observed upregulation of *Cdkn1a* in both intestine and tongue and a marked decrease in the cell proliferation marker KI67 soon after commencing 5-FU administration. In tongue tissues, the upregulation of *Cdkn1a* was temporary occurring in the early period after starting 5-FU, while the intestine showed sustained upregulation. Although our study is not mechanistic, it suggests that control of the cell cycle though the action of p53 is the key event leading to the atrophy of the oral and intestinal epithelium, which is the main cellular compartment affected by 5-FU. Furthermore, other genes not related to *Cdkn1a* but also involved in cell cycle regulation were strongly upregulated in response to 5-FU. These included *Ccng1* (cyclin G1), which has been shown to act independently of *Cdkn1a* to induce cell cycle arrest in response to DNA damage^56^ and was found here as upregulated in the intestine. Another cell cycle regulator, *Gtse1* (G2 and S phase expressed protein), showed one of the highest increases in fold change in both tongue and intestine. *Gtse1* has been shown to be involved in delaying the G2/M phase of the cell cycle and to negatively impact p53 activity, perhaps acting as a feedback loop regulator^57^. *Gtse1* has also been shown to suppress apoptosis and mediate resistance to the chemotherapeutic cisplatin^58^. The roles of different p53 targets in mediating in vivo responses to DNA damage are not fully understood and therefore further studies are warranted to dissect which genes involved in cell cycle regulation are critical mediators of epithelial atrophy and its recovery following chemotherapy.

Although cell cycle arrest was the dominant phenotypic response to 5-FU, we observed transcriptional upregulation of both cell cycle checkpoint regulators and pro-apoptotic genes, even within single cells as revealed through scRNASeq of tongue mucosa. These data suggest that p53 does not control cell fate by favoring a specific transcriptional program within cells. These results are consistent with a model in which p53 unselectively binds to its target genes independent of their biological function, with the decision of whether a cell undergoes cell cycle arrest or apoptosis determined by other factors such as the balance between pro-apoptotic, anti-apoptotic and anti-proliferative mediators within cells^59^. In this respect, we observed here that effectors from the anti-apoptotic *Bcl2* gene family were upregulated within the epithelial compartment after 5-FU treatment with protein co-expression of BCL2 and BAX detected within cells. Anti-apoptotic BCL2 proteins protect cells from stress-induced death by inhibiting BAX/BAK-mediated mitochondrial permeabilization and suppressing activation of the intrinsic apoptotic pathway^60^. Further supporting the idea that the ratios of anti-proliferative and pro-apoptotic proteins is important to determine cell fate, immunostaining showed a dramatic upregulation of CDKN1a in 5-FU-treated animals compared to controls, whereas differences in levels of the pro-apoptotic protein BAX between homeostasis and after 5-FU treatment were less dramatic. Altogether, our study provides in vivo evidence that helps to further understanding of the manner in which the transcriptional factor p53 operates in vivo within cells.

The transcriptional program activated by p53 in oral and intestinal tissues showed similarities and also distinct p53 target genes differentially expressed at each site. These results agree with previous work that showed the p53-mediated induction of downstream targets varies across tissues^18,20^, and highlight that although the oral and small intestine are part of the same organ, behavior and responses in these barriers are specific and thus, there is a need to understand injury in each of these compartments to develop targeted approaches against chemotherapy toxicity. Eight universal targets of p53 across tissues were previously identified in a study that did not include oral tissues^20^. Here we found that seven out these eight genes were indeed upregulated in both tongue and intestine in response to 5-FU. *Ccng1* was the only universal target gene not showing upregulation in tongue in our study, although it showed a statistically significant p value and increased expression in 5-FU-treated animals compared to PBS, but the change was below the one log_2_ fold threshold employed to define significant genes. However, several p53 target genes showed site specificity. For instance, the pro-apoptotic mediators *Fas* and *Pmaip1* (Noxa) showed upregulation in tongue but not in intestine. In agreement with the mouse model employed here, we have previously shown upregulation of *Pmaip1* in human oral epithelial cells collected from patients undergoing chemotherapy^3^. Moreover, outside the p53 pathway, we found that oral, but not intestinal tissues upregulated genes involved in fatty acid uptake and oxidation, suggestive of metabolic reprogramming, while this response was not observed in the intestine. Upregulation of lipid metabolism-related genes in tongue occurred mostly at day 4 of 5-FU administration coinciding with cell cycle re-entry. Fatty acid uptake and oxidation generate considerably more energy per carbon than glucose, and have been shown to mediate resistance of cancer cells to the chemotherapeutics cisplatin^61^. Enhanced fatty-acid uptake and oxidation in the oral epithelium may act as a bioenergetic bridge from cytostasis to regeneration, ensuring that proliferative recovery is coupled to sufficient energy supply.

Our work mapped cellular populations and their kinetics in response to 5-FU in the tongue. While the progenitor populations responsible for renewal of the intestinal epithelium under homeostasis and in response to injury are well characterized^62,63^, there is a paucity of information on the defining features of oral epithelial cell progenitors. A previous study performed a scRNASeq analysis of β4-integrin-sorted epithelial cells from the buccal mucosa of mice, finding that the basal cell layer contains progenitors and post-mitotic cells at various stages of maturation^64^. Agreeing with these results, here we observe cells with basal identity but representing different states including quiescent, moderately proliferative and highly proliferative. These populations showed differences in their sensitivity to 5-FU with highly proliferative cells becoming severely depleted soon after 5-FU administration, while moderately proliferative cells were more resistant. Among quiescent cells, a population of putative oral epithelial progenitors enriched for genes involved in Wnt-signaling was identified in homeostasis with its proportions decreasing during chemotherapy and returning to homeostatic levels after 5-FU withdrawal. These chemosensitive cells may represent progenitors charged with maintenance of homeostasis but incapable of surviving chemotoxic stress. In contrast, we also identified a population of putative oral epithelial cell progenitors with a p53 and AP-1 complex gene signature that showed resistance to 5-FU and appeared to expand at day 4, when cell cycle re-entry occurred. In the lung, progenitors with similar gene signatures have been shown to act as transitional cells responsible for tissue regeneration after injury^21–23^. Intestinal tissues have also been shown to harbor a pool of progenitors in charge of tissue renewal under homeostasis^65^, while a distinct population of transitional progenitor cells emerges under injury and is necessary for tissue reconstitution after irradiation^62^. Contrary to the lung and intestine, however, the putative oral progenitors carrying a transitional-like gene program enriched for p53 and AP-1 complex signatures revealed by our study were seen to exist under homeostasis, rather than being transitional cells induced after injury. The dynamics of the identified populations and their regenerative capacity in the tongue epithelium after chemotoxic stress warrant further mechanistic studies.

In conclusion, our work provides a deep phenotypic characterization of the responses in oral and intestinal tissues to the chemotherapeutic 5-FU showing that mucositis is a non-inflammatory condition in which disruption in the replication of epithelial progenitor populations leads to tissue atrophy and loss of mucosal integrity. By mapping gene expression dynamics, we also show that p53 is the master regulator of responses to 5-FU in both compartments with concomitant induction within tissues and cells of transcriptional programs controlling divergent cell fates but with cell cycle arrest as the dominant phenotypic response. Through focused scRNASeq of the oral mucosa we map the kinetics of cell populations in response to 5-FU, revealing novel populations of putative oral epithelial cell progenitors with different susceptibilities to chemotoxic stress. A population with an AP-1 complex and p53-driven stress-adaptive signature showed resistance to 5-FU emerging as a candidate population that mediates tissue renewal after 5-FU treatment.

## Acknowledgements

This study was funded through grants R56 DE028545 and R01 DE032131 from the National Institutes of Health. Lu Li was supported by grant K99 DE 034829 from the National Institutes of Health.

