## Supplementary Figures for "Chemotherapy-Induced Oral Mucosal Injury Is Defined by p53 Activation, Cell Cycle Arrest and Diverse Epithelial Progenitor Dynamics"

**Supplementary Figure S1. Systemic 5-fluorouracil induces progressive mouse weight loss and epithelial clefting of the tongue dorsal mucosa.** (A) Longitudinal change in body weight in animals treated with 5-fluorouracil (5-FU) or PBS over a 10-day observation period. Body weight is shown as percentage change from baseline. Data are presented as mean ± SD. (B) Representative hematoxylin and eosin (H&E)-stained sections of tongue dorsal mucosa showing epithelial clefting after 5-FU treatment. Scale bars = 50μm.

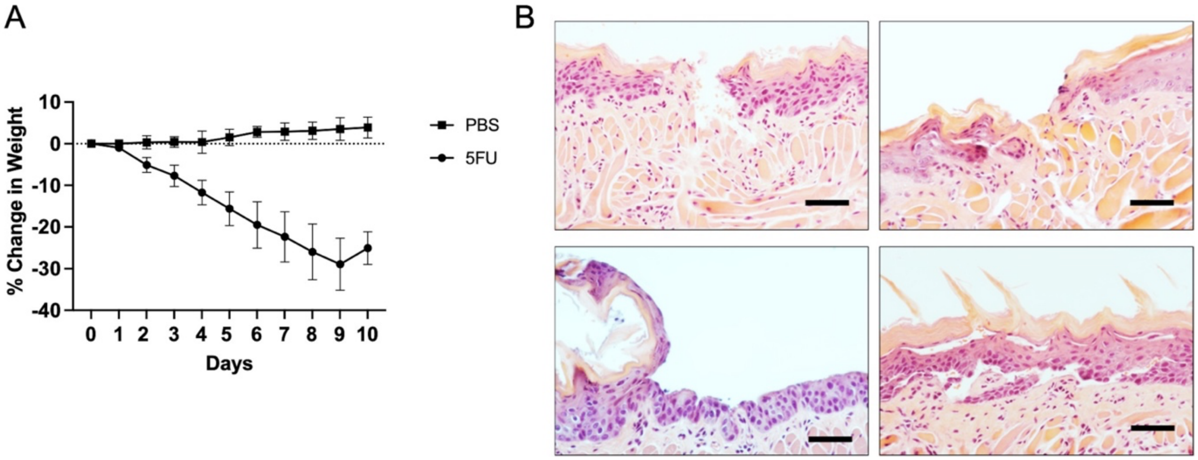

**Supplementary Figure S2. Clinical presentation of oral mucositis lesions in patients undergoing cancer treatment.** Clinical signs of mucositis were monitored at a baseline visit, prior to commencing chemotherapy, and at periodic visits as indicated for up to 42 days. Images illustrate representative therapy-associated alterations of dorsal tongue papillary architecture and oral mucosal integrity. (A) Patient undergoing chemotherapy for breast cancer. At baseline, the dorsal tongue shows preserved filiform papillae with the characteristic "brush-like" architecture. At Day +9, patchy erythema and progressive attenuation of filiform papillae are observed. By Day +14, the dorsal tongue demonstrates marked loss of filiform papillae, resulting in a smoother, glossier surface. At Day +28, the patient presented with a lesion in lip commissure consistent with angular cheilitis due to mucositis. (B) Patient undergoing chemotherapy for gastrointestinal (rectum) cancer. Baseline (Day 0) image shows the dorsal tongue with intact papillary architecture and the typical "brush-like" pattern of filiform papillae. At Day +42, the dorsal tongue appears noticeably smoother, reflecting partial loss and flattening of filiform papillae. Additional views at the same time point show involvement of the left and right labial commissures, with erythema and lesions consistent with angular cheilitis due to mucositis. (C) Patient treated with chemotherapy and radiation for head and neck cancer. At baseline (Day 0), the dorsal tongue exhibits well-defined filiform papillae and the typical "brush-like" dorsal surface morphology. By Day +31, the dorsal tongue shows pronounced loss of papillae, erythema and an ulcer-like lesion. Additional images demonstrate involvement of the left lip and labial commissure.

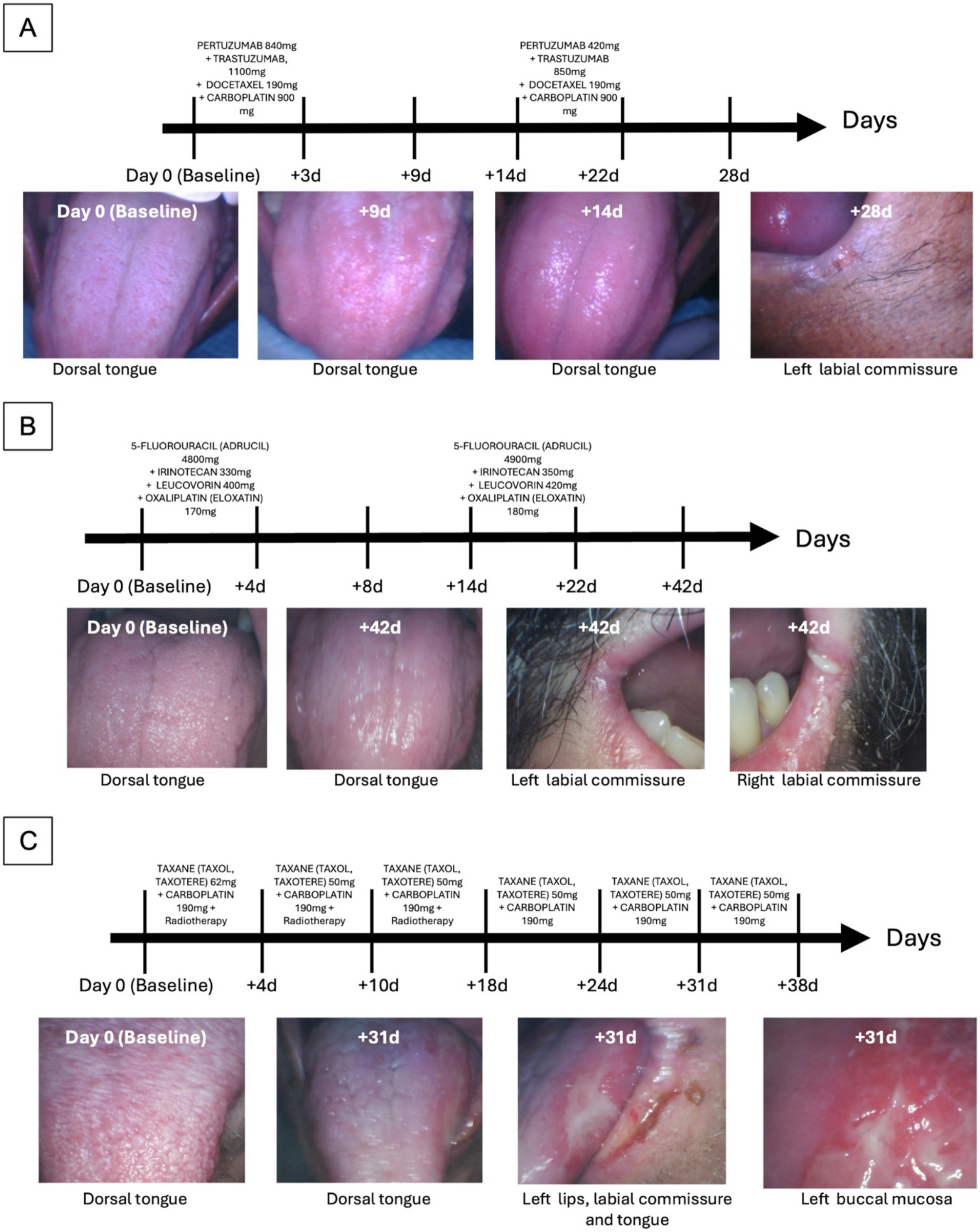

A
PERTUZUMAB 840mg + TRASTUZUMAB, 1100mg + DOCETAXEL 190mg + CARBOPLATIN 900 mg
PERTUZUMAB 420mg + TRASTUZUMAB 850mg + DOCETAXEL 190mg + CARBOPLATIN 900 mg
Days
Day 0 (Baseline)
+3d
+9d
+14d
+22d
28d
Day 0 (Baseline)
+9d
+14d
+28d
Dorsal tongue
Dorsal tongue
Dorsal tongue
Left labial commissure
B
5-FLUOROURACIL (ADRUCIL) 4800mg + IRINOTECAN 330mg + LEUCOVORIN 400mg + OXALIPLATIN (ELOXATIN) 170mg
5-FLUOROURACIL (ADRUCIL) 4900mg + IRINOTECAN 350mg + LEUCOVORIN 420mg + OXALIPLATIN (ELOXATIN) 180mg
Days
Day 0 (Baseline)
+4d
+8d
+14d
+22d
+42d
Day 0 (Baseline)
+42d
+42d
+42d
Dorsal tongue
Dorsal tongue
Left labial commissure
Right labial commissure
C
TAXANE (TAXOL, TAXOTERE) 62mg + CARBOPLATIN 190mg + Radiotherapy
TAXANE (TAXOL, TAXOTERE) 50mg + CARBOPLATIN 190mg + Radiotherapy
TAXANE (TAXOL, TAXOTERE) 50mg + CARBOPLATIN 190mg + Radiotherapy
TAXANE (TAXOL, TAXOTERE) 50mg + CARBOPLATIN 190mg
TAXANE (TAXOL, TAXOTERE) 50mg + CARBOPLATIN 190mg
TAXANE (TAXOL, TAXOTERE) 50mg + CARBOPLATIN 190mg
Days
Day 0 (Baseline)
+4d
+10d
+18d
+24d
+31d
+38d
Day 0 (Baseline)
+31d
+31d
+31d
Dorsal tongue
Dorsal tongue
Left lips, labial commissure and tongue
Left buccal mucosa

**Supplementary Figure S3. Temporal dynamics of CD45+ immune cell infiltration in tongue and intestinal mucosa following 5-fluorouracil treatment.** (A, C) Representative images of dorsal tongue (A) and small intestinal mucosa (C) tissue sections after immunohistochemical staining for CD45 (n = 4 mice per condition). CD45+ leukocytes are visualized as brown chromogenic staining, while tissue architecture and nuclei are visualized by hematoxylin counterstaining (blue). A positive control (PC) section of mouse spleen tissue confirms robust CD45 immunoreactivity. Scale bars= 100µm. (B, D) Quantification of CD45+ immune cells in tongue (B) and in intestinal mucosa (D). Individual points represent biological replicates, and bars indicate mean ± SD. *p<0.05, **p<0.01, as determined via ANOVA with Tukey post hoc analysis.

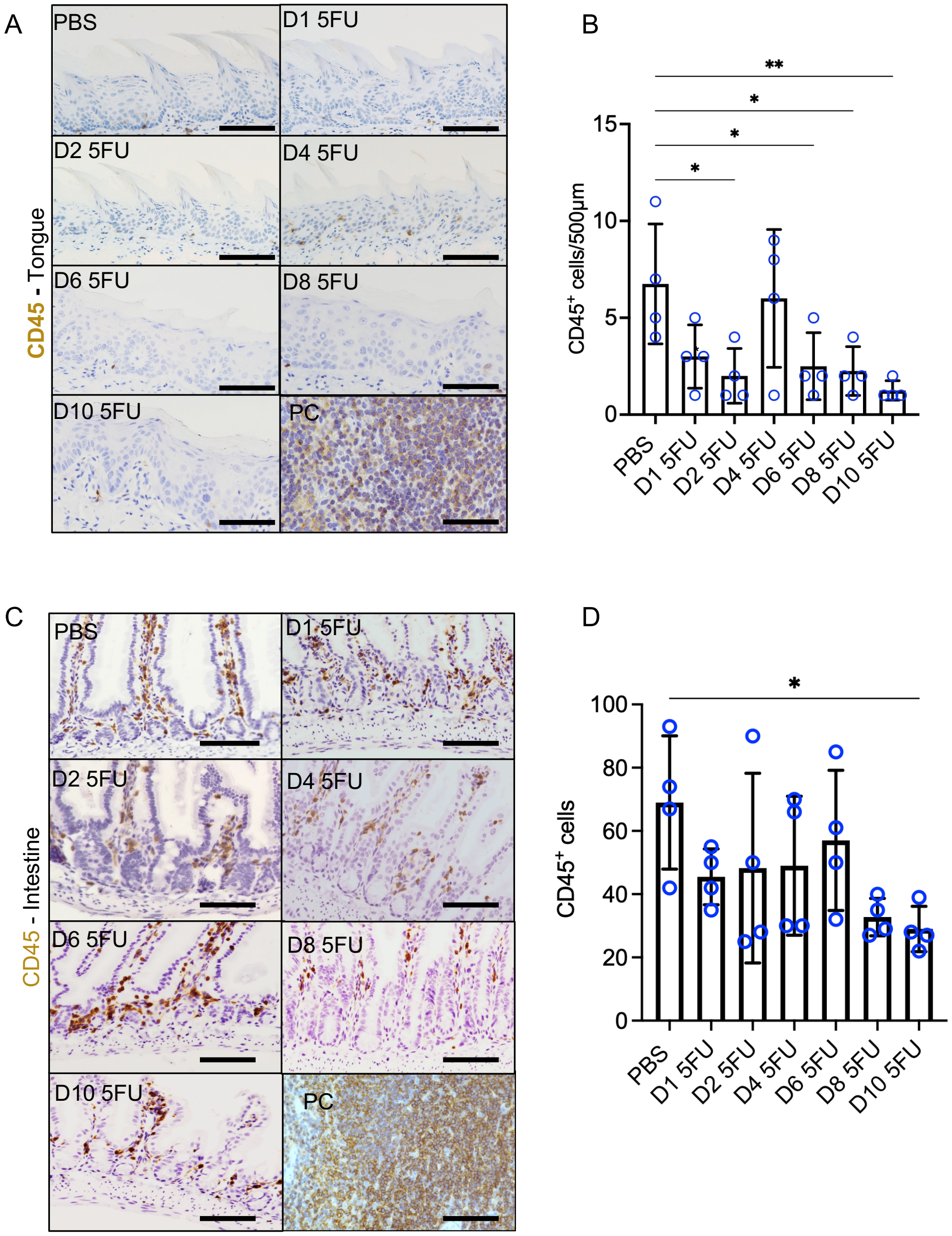

**Supplementary Figure S4. Limited apoptotic activity in tongue and intestinal mucosa following 5-fluorouracil treatment.** (A-D) Representative immunohistochemical staining and quantification for cleaved Caspase-3 in tongue (A-B) and small intestine (C-D) following 5-FU administration (n= 4 mice per condition). Cleaved caspase-3–positive cells appear brown, while nuclei are visualized by hematoxylin counterstaining (blue). A positive control (PC) section of mouse subcutaneous tumor confirms antibody specificity. No significant differences found via ANOVA. (E-F) TUNEL staining of tongue sections across the same experimental timeline. Apoptotic nuclei are detected as green fluorescence, while nuclei counterstained with DAPI are blue. A tongue tissue section treated with DNAse I served as PC. Scale bars = 100 µm. *p<0.05 as determined via ANOVA with Tukey post hoc analysis.

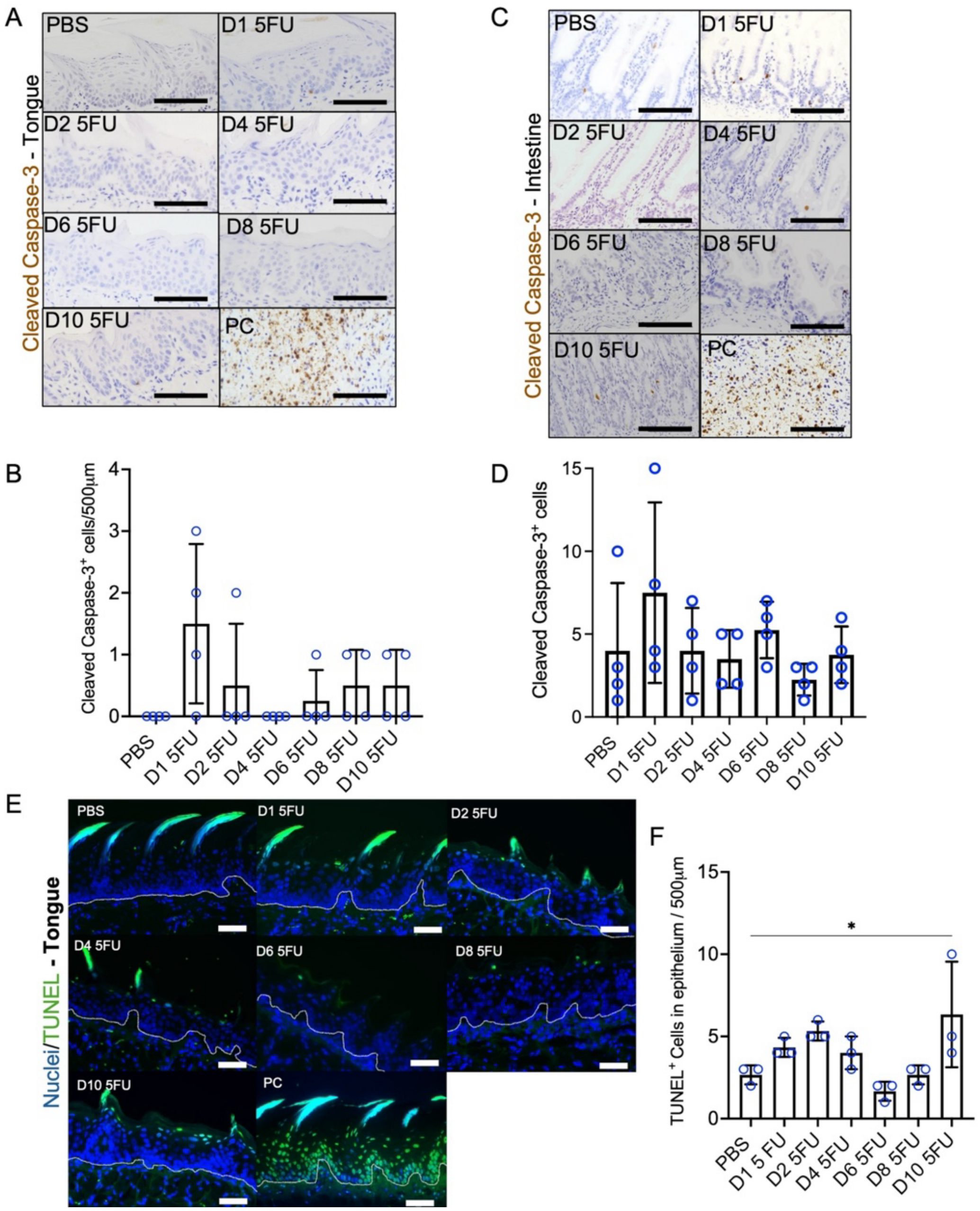

**Supplementary Figure S5. Main cell types identified via single-cell RNASeq of tongue and their change in proportions following 5-FU treatment.** (A) Heatmap showing cell-type-defining marker genes and their expression in homeostasis. Rows represent marker genes and columns represent individual cells grouped by annotated cell populations. (B) UMAP visualization of cell types in tongue and their change in abundance following 5-FU treatment. Each point represents a single cell annotated via module scores calculated using cell-type-defining marker genes.

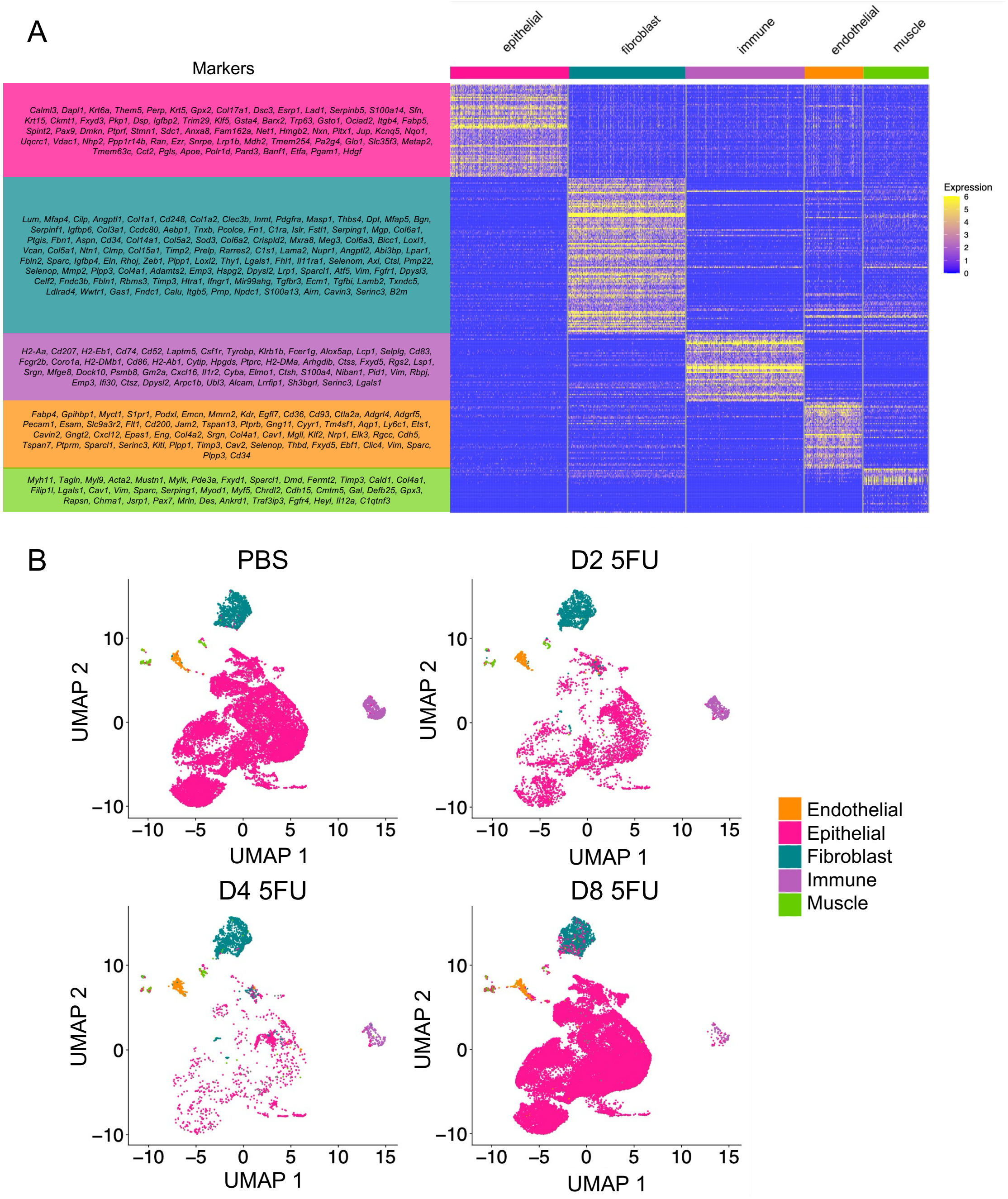

**Supplementary Figure S6. Epithelial subclusters.** (A) UMAP projection of scRNASeq data from tongue in PBS- and 5-FU-treated control animals showing 22 epithelial subclusters. (B) Heatmap and hierarchical clustering of 716 marker genes identified after each epithelial cluster in the PBS condition was compared to non-epithelial cells. Hierarchical clustering reveals transcriptional relationships among epithelial clusters separating suprabasal, basal and salivary epithelial populations.

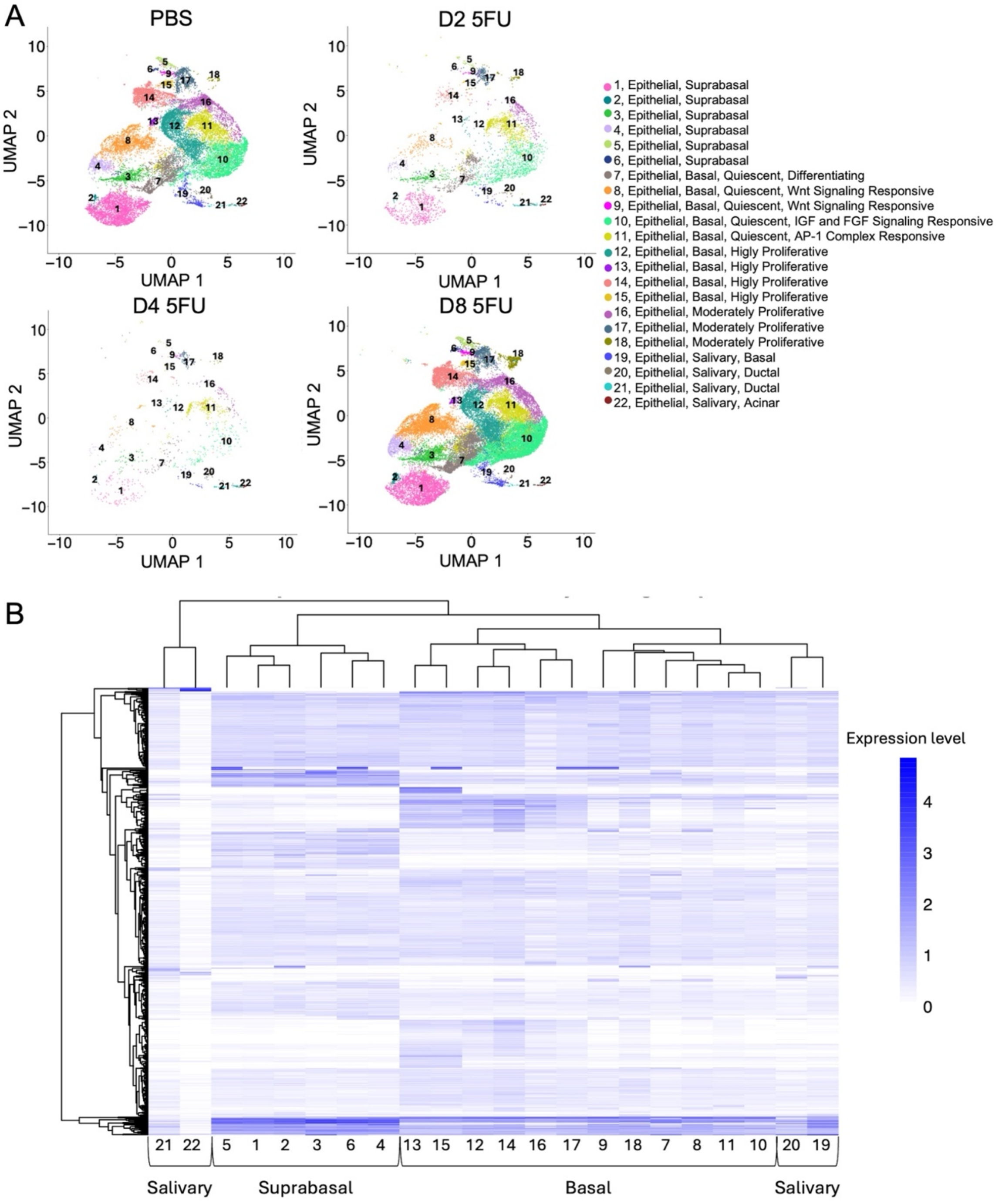

**Supplementary Figure S7. Temporal shifts in basal and suprabasal epithelial populations following 5-FU treatment.** (A) UMAP projection of epithelial cells from tongue dorsum PBS-a and 5-FU-treated animals. Each point represents a single epithelial cell annotated and colored via module scores calculated using basal- and suprabasal-defining marker genes. Salivary gland cells were exclude from these graphs.

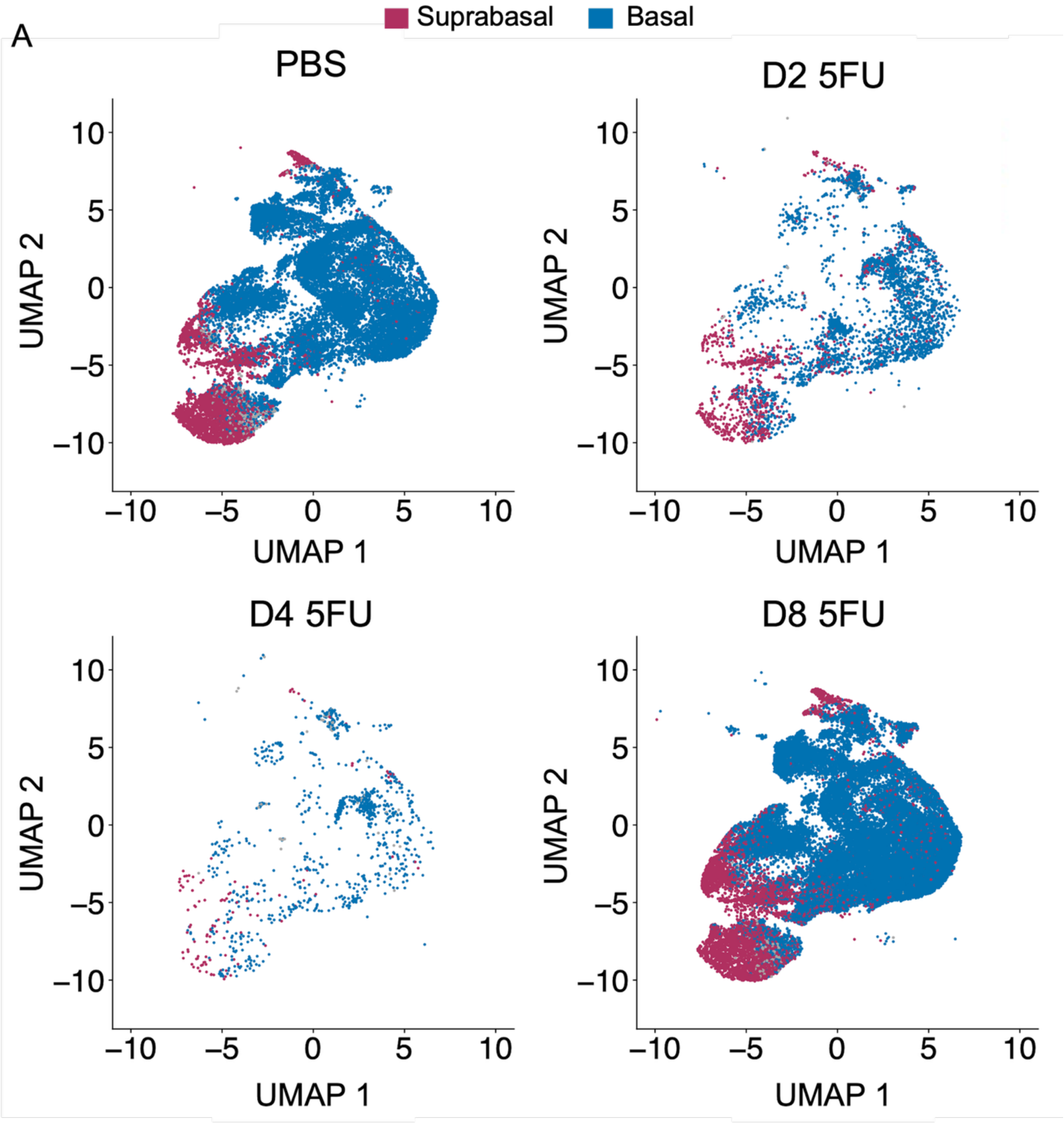

**Supplementary Figure S8. Changes in abundance of tongue epithelial subpopulations following 5-FU treatment a revealed by scRNASeq.** (A) UMAP visualization of epithelial cells in PBS- and 5-FU-treated mice. Each point represents a single epithelial cell annotated and colored via module scores calculated using epithelial subpopulation marker genes. (B) Heatmap showing the relative proportion of epithelial subclusters across time points.

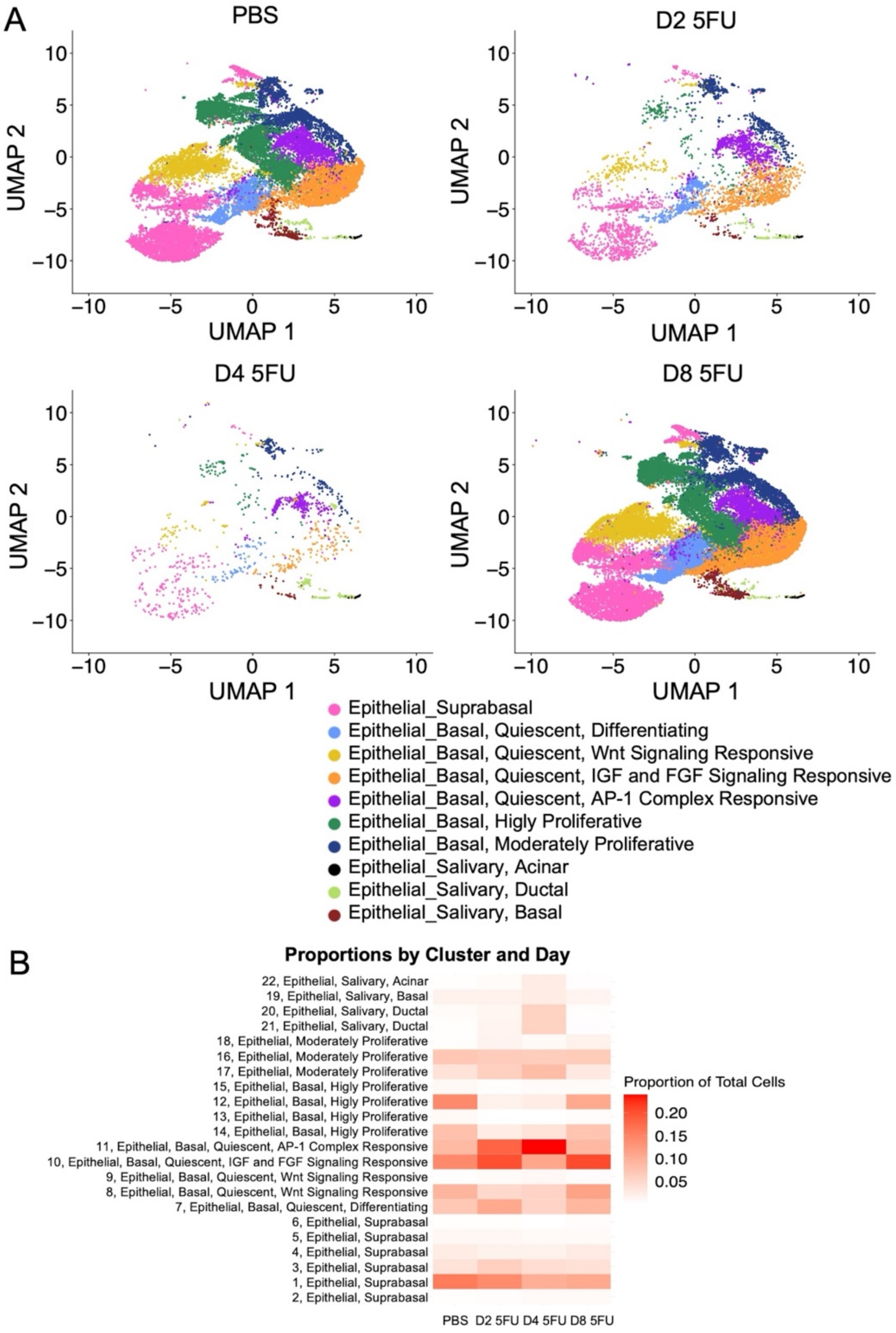

**Supplementary Figure S9. EpCAM identifies minor salivary glands in tongue.** (A) Representative immunofluorescence image of the tongue dorsum epithelium and (B) Representative immunofluorescence image of tongue salivary gland tissue stained for EpCAM (green). Scale bars, 50 µm.

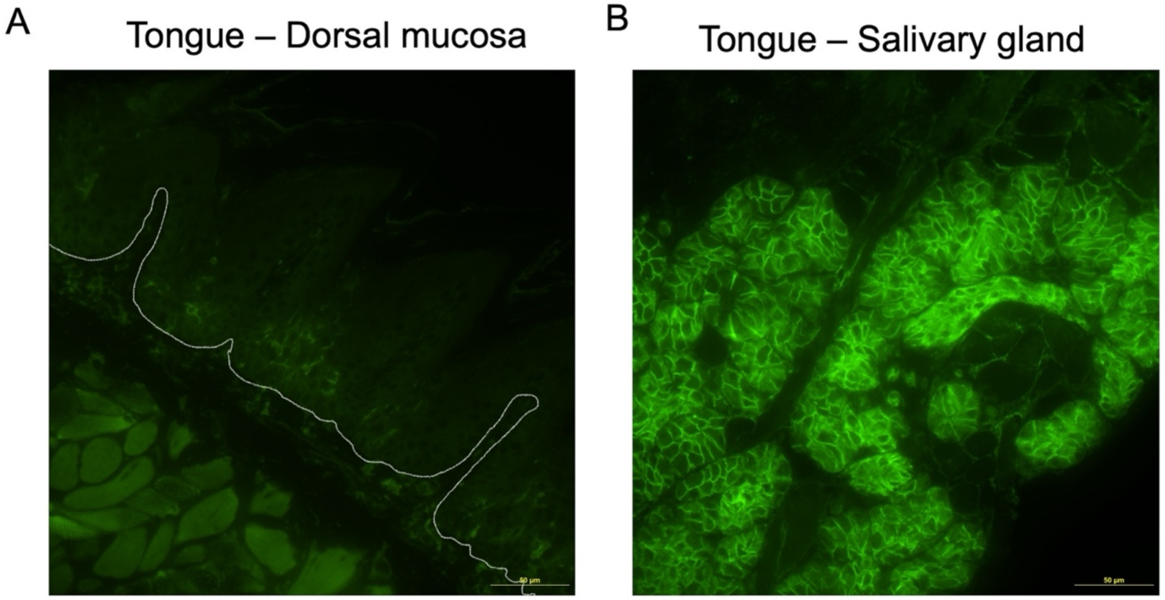

**Supplementary Figure S10 - Pathway enrichment analysis across stromal and immune cell populations following 5-FU treatment.** Gene Ontology (GO) and KEGG pathway (K) enrichment analyses were performed on differentially expressed genes derived from sc RNASeq data.

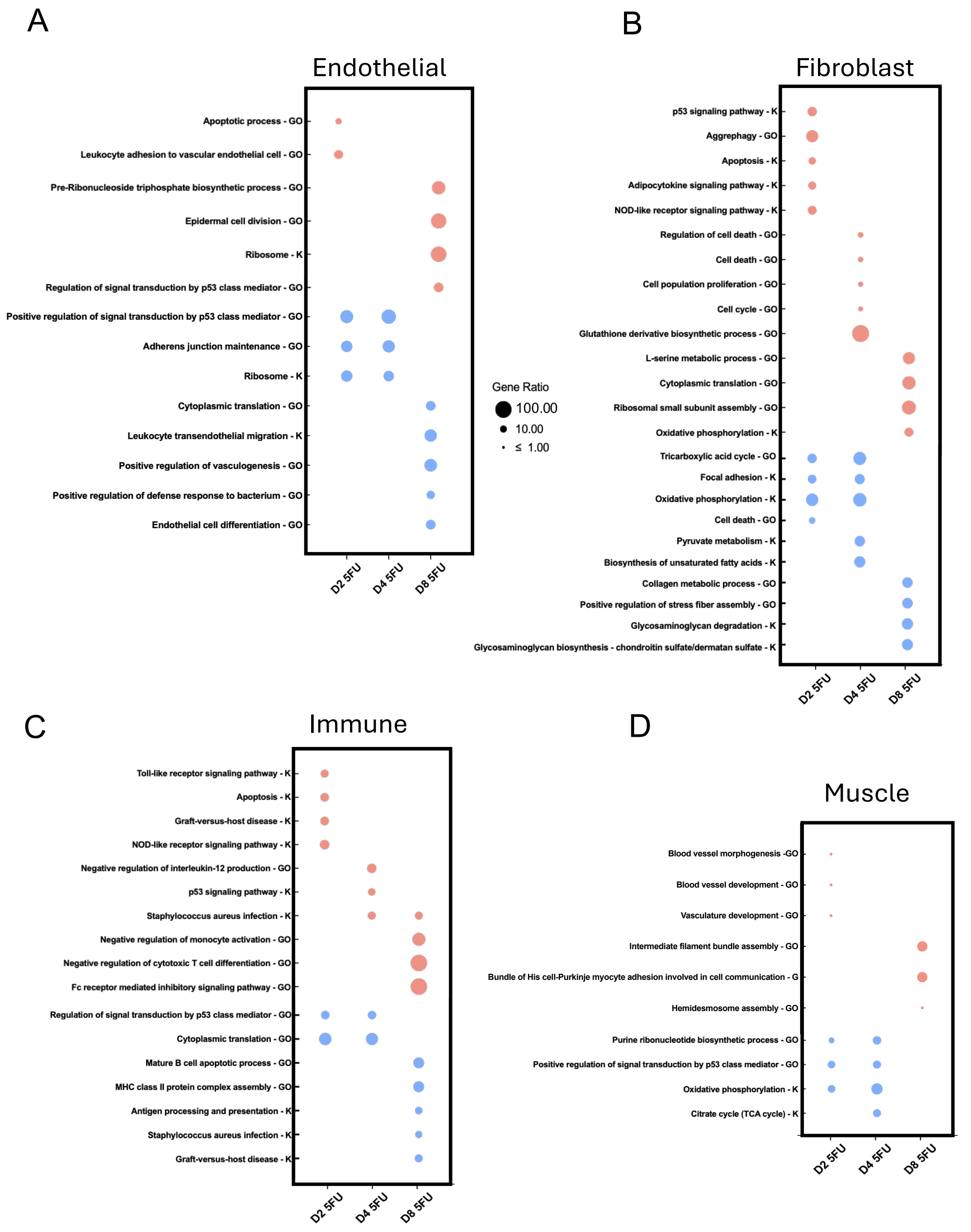

**Supplementary Figure S11 - Single-cell co-expression analysis of Cdkn1a (p21) with apoptosis- and stress-related genes in epithelial basal and suprabasal populations following 5-FU treatment.** Heatmaps display Cdkn1a expression on the x-axis and the indicated gene on the y-axis, with color intensity representing the percentage of cells within each expression bin. (A) Co-expression of Cdkn1a and Bid showing that Bid expression is detected in a subset of epithelial cells and tends to occur in cells with higher Cdkn1a expression. (B) Co-expression of Cdkn1a and Bik, where Bik-expressing cells largely overlap with Cdkn1a-positive epithelial cells. (C) Co-expression of Cdkn1a and Bbc3 (PUMA) demonstrating that when Bbc3 expression is detected, it predominantly occurs in Cdkn1a-expressing cells. (D) Co-expression of Cdkn1a and Pmaip1 (NOXA) showing a similar pattern in which Pmaip1-positive cells largely coincide with Cdkn1a-high epithelial cells. (E) Co-expression of Cdkn1a and Casp3, indicating that Casp3 expression is observed in a subset of epithelial cells and overlaps with Cdkn1a expression. (F) Co-expression of Cdkn1a and Fas, where Fas-expressing epithelial cells are predominantly Cdkn1a-positive. (G) Co-expression of Cdkn1a and Eda2r showing partial overlap between Eda2r expression and Cdkn1a-positive epithelial cells. (H) Co-expression of Cdkn1a and Tnfsf12 (TWEAK) demonstrating detectable Tnfsf12 expression in subsets of epithelial cells with overlapping Cdkn1a expression. (I) Co-expression of Cdkn1a and Tnfrsf18 (GITR) indicating that Tnfrsf18 expression occurs primarily in epithelial cells that also express Cdkn1a.

A

#### Co-expression *Cdkn1a* and *Bid*

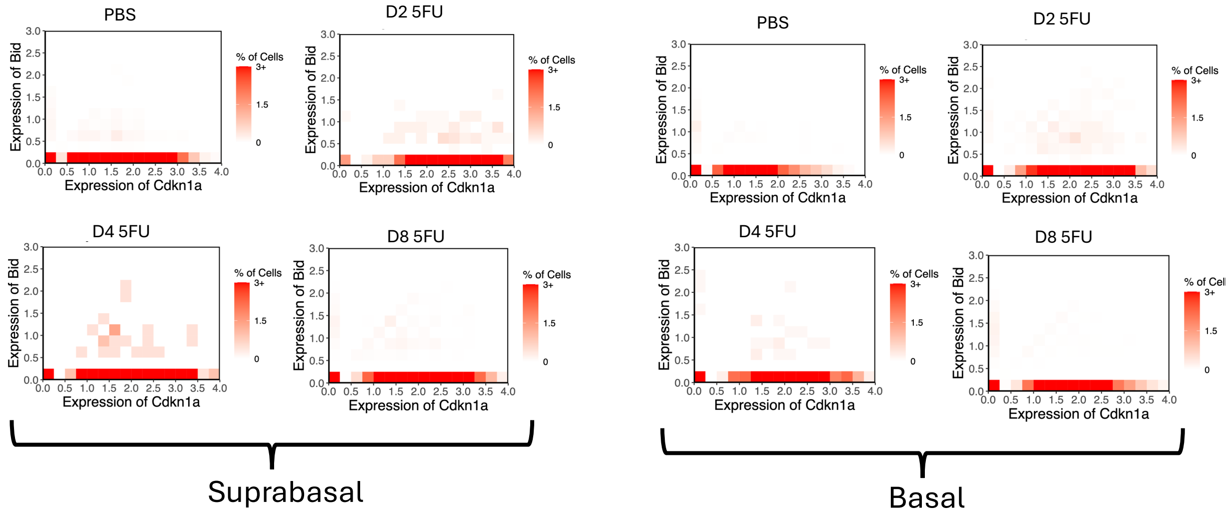


B

#### Co-expression *Cdkn1a* and *Bik*

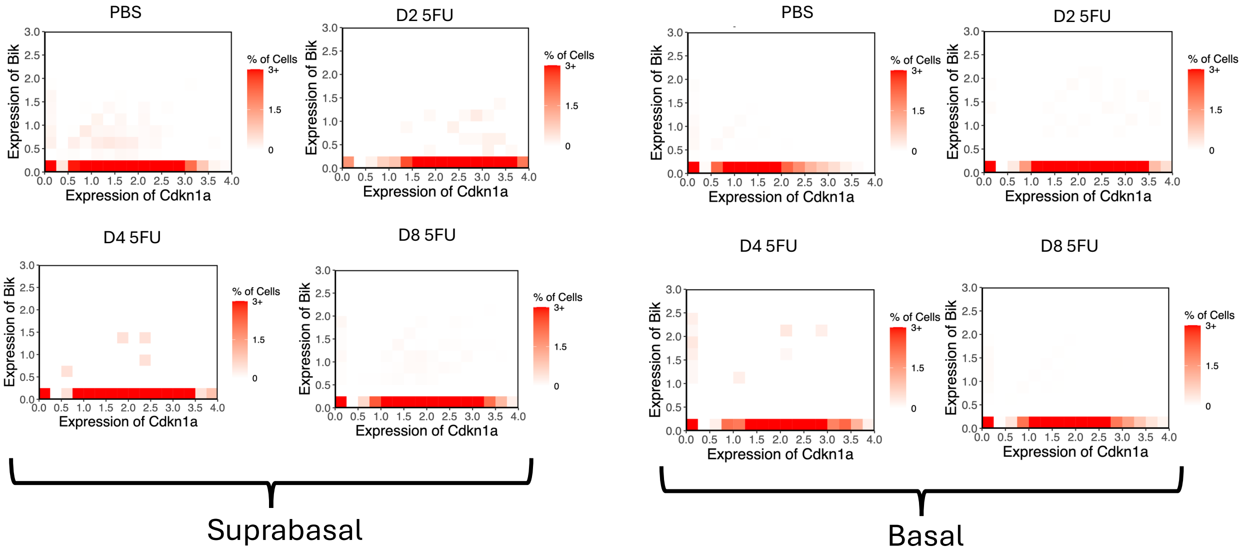

C

#### Co-expression *Cdkn1a* and *Bbc3*

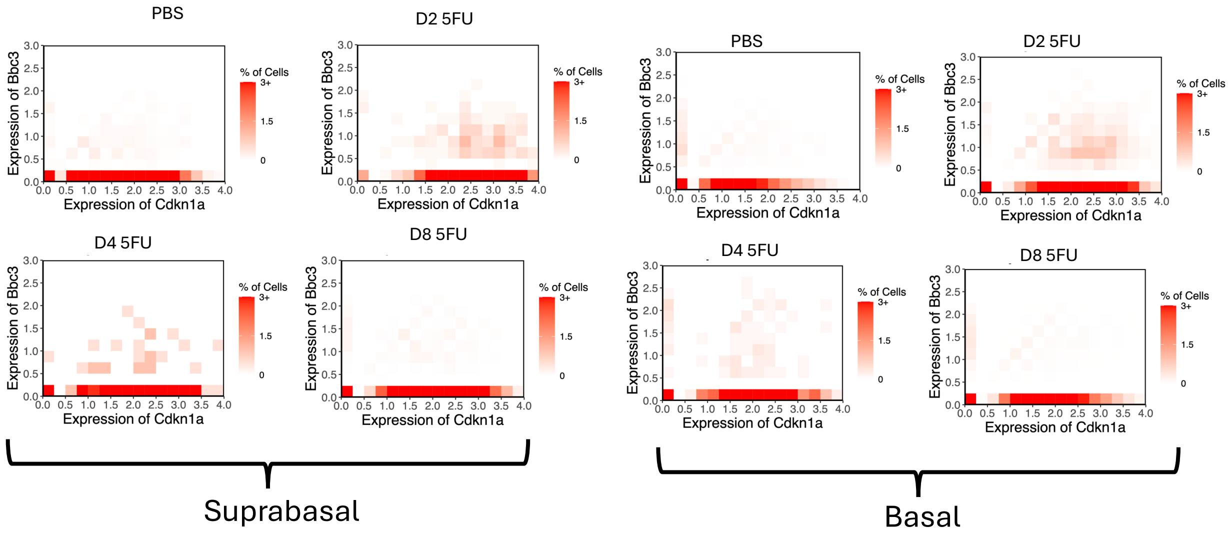


D

#### Co-expression *Cdkn1a* and *Pmaip1*

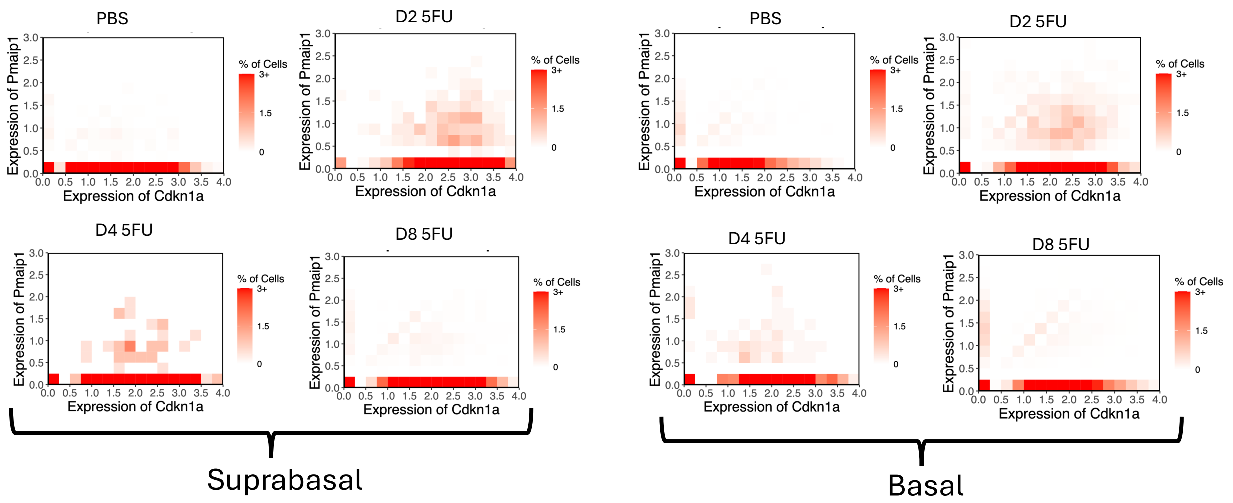

E

### Co-expression *Cdkn1a* and *Casp3*

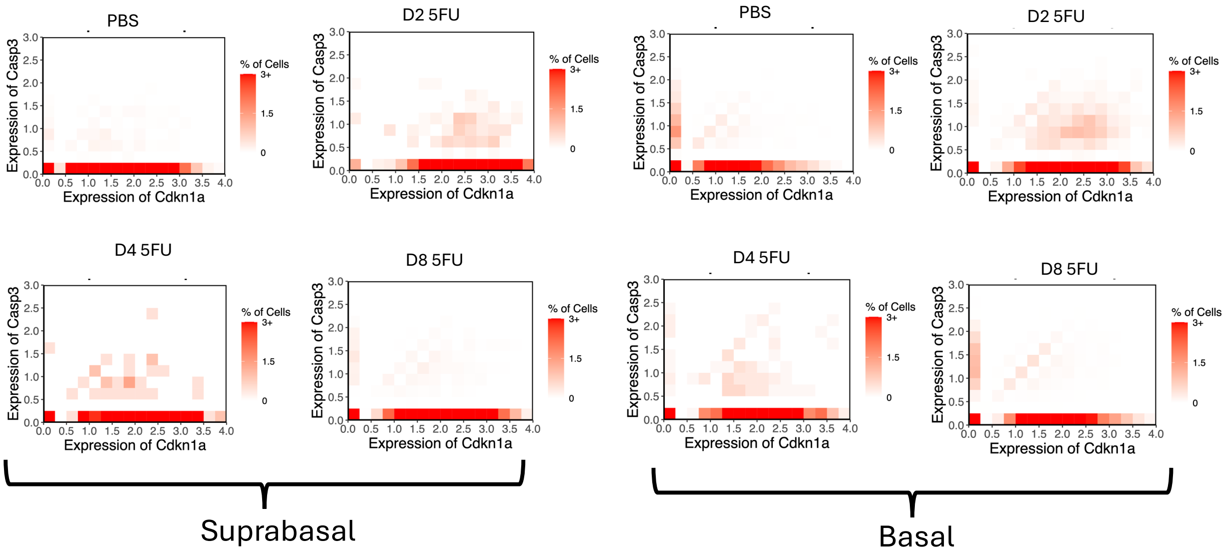


F

### Co-expression *Cdkn1a* and *Fas*

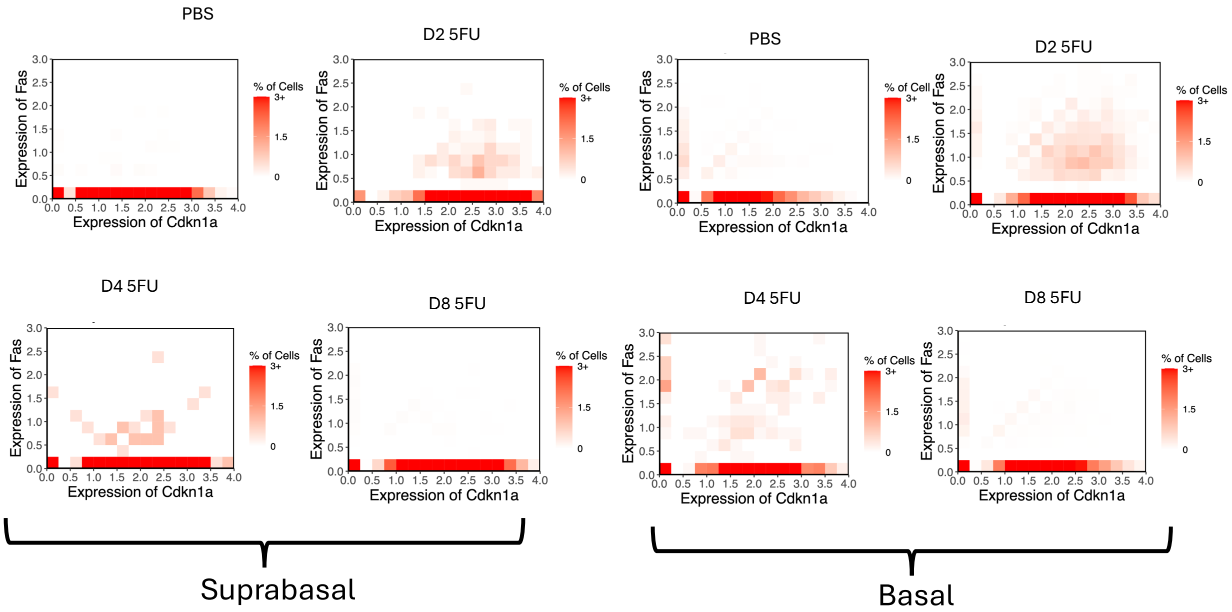

G

#### Co-expression *Cdkn1a* and *Eda2r*

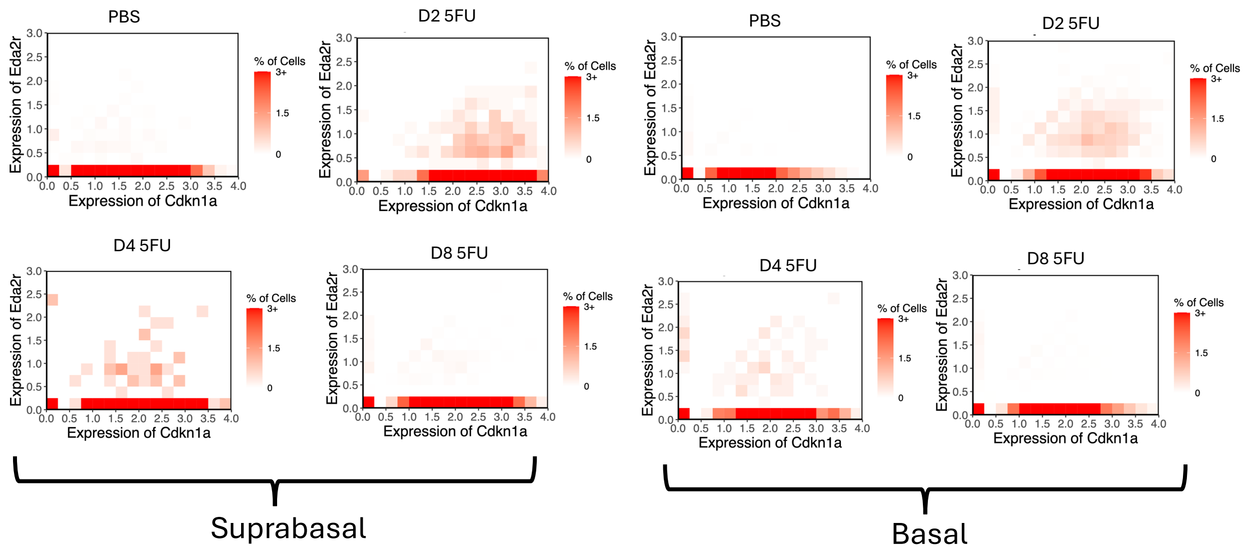


H

#### Co-expression *Cdkn1a* and *Tnfrsf12a*

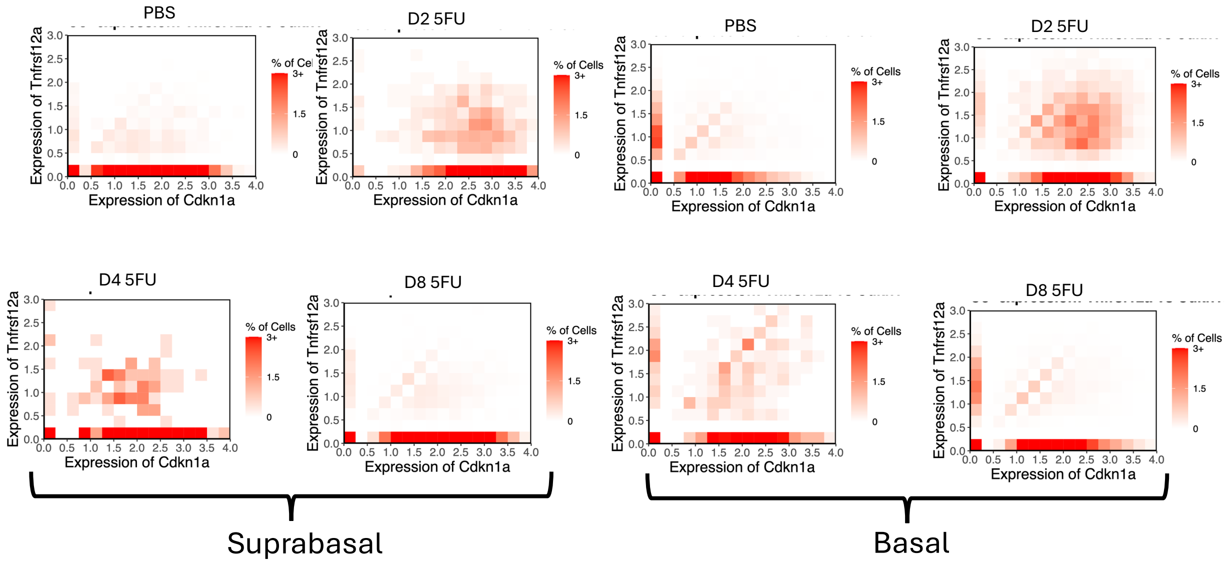

I

### Co-expression *Cdkn1a* and *Tnfrsf18*

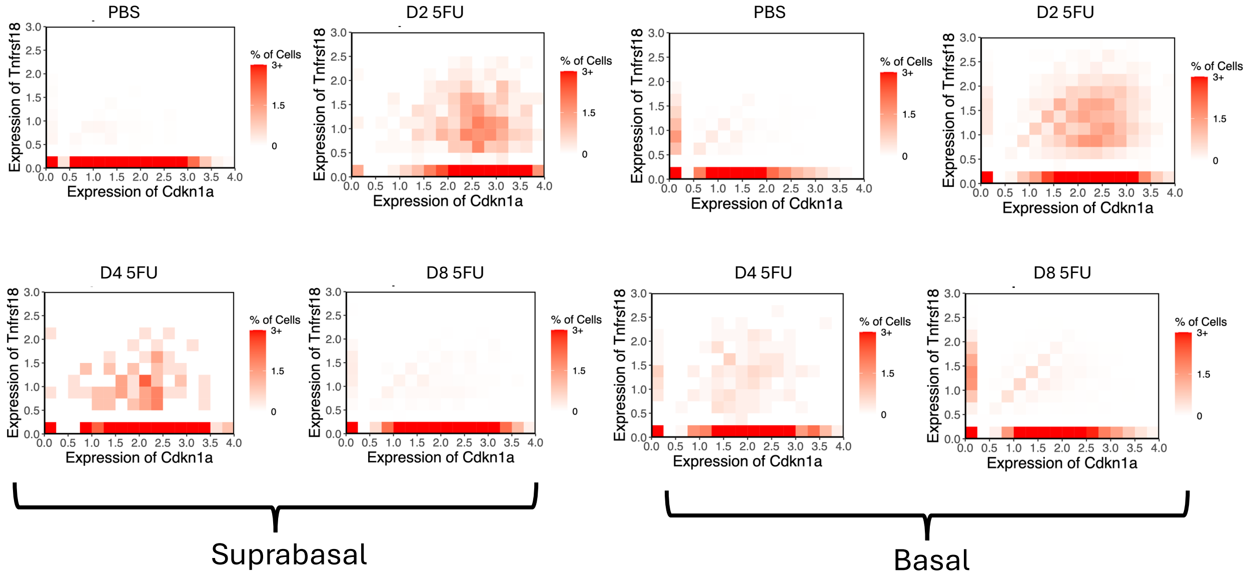
